# Integron activity accelerates the evolution of antibiotic resistance

**DOI:** 10.1101/2020.08.07.237602

**Authors:** Célia Souque, José A. Escudero, R.Craig MacLean

**Affiliations:** University of Oxford, Department of Zoology, 11a Mansfield Road, Oxford, OX1 3ZY; Universidad Complutense de Madrid, Departamento de Sanidad Animal, Avenida Puerta de Hierro, s/n., 28040 Madrid

## Abstract

Mobile integrons are widespread genetic platforms that allow bacteria to modulate the expression of antibiotic resistance cassettes by shuffling their position from a common promoter. Antibiotic stress induces the expression of an integrase that excises and integrates cassettes, and this unique recombination and expression system is thought to allow bacteria to ‘evolve on demand’ in response to antibiotic pressure. To test this hypothesis, we inserted a custom three cassette integron into *P. aeruginosa*, and used experimental evolution to measure the impact of integrase activity on adaptation to gentamicin. Crucially, integrase activity accelerated evolution by increasing the expression of a gentamicin resistance cassette through duplications and by eliminating redundant cassettes. Importantly, we found no evidence of deleterious off-target effects of integrase activity. In summary, integrons accelerate resistance evolution by rapidly generating combinatorial variation in cassette composition while maintaining genomic integrity.

## Introduction

Given the mounting threat posed by antibiotic resistance we need a better understanding of the mechanisms used by bacteria to evolve resistance to antibiotics. Mobile integrons (MIs) are widespread elements providing a platform for the acquisition, shuffling, and expression of gene cassettes, many of which are antibiotic resistance genes (Recchia & Hall, 1995; Escudero et al., 2015). These elements are typically associated with transposons and conjugative plasmids, and have played an important role in the evolution of resistance in pathogenic bacteria (Partridge et al., 2018). Five classes of MIs have been described but the class 1 integron is, by far, the most prevalent and clinically relevant. The first multidrug resistance plasmids that were isolated in the 1950s carried class 1 mobile integrons (Liebert et al., 1999; Mitsuhashi et al., 1961; Rownd et al., 1966; Stokes & Hall, 1989), and recent surveys have shown that class 1 integrons are found in a substantial fraction of isolates of *E. coli* (Halaji et al., 2020; Rao et al., 2006; Yu et al., 2003), *K. pneumoniae* (Firoozeh et al., 2019; Li et al., 2013), *P. aeruginosa* (Oliver et al., 2015; Ruiz-Martínez et al., 2011), and *A. baumanii* (Chen et al., 2015; Turton et al., 2005).

Mobile integrons consist of an integrase encoding gene named *intI* followed by a recombination site, *attI* (R M Hall et al., 1991; Ruth M. Hall et al., 2000) and a variable array of mobile gene cassettes (typically 2-5) ending each in a characteristic hairpin recombination site called the *attC* site (R M Hall et al., 1991). Integron cassettes usually lack a promoter, and their expression is driven by the Pc promoter located upstream of the array (Collis & Hall, 1995), such that expression levels are highest for cassettes closest to the promoter (Collis & Hall, 1995). The SOS response induces the expression of the integrases (Guerin et al., 2009), which then allows for the efficient integration and excision of cassettes in the array through *attC × attI* and *attC × attC* reactions, respectively (Collis & Hall, 1992). A peculiarity of this system is that integron recombination is semi-conservative, as only the bottom strand of the cassette is excised from the array through recombination events which include a replication step (Loot et al., 2012). The implication of this is that cassette excision and re-integration can be assimilated to either a ‘cut and paste’ process, resulting in the movement of a cassette within an array, or to a ‘copy and paste’ process, leading to the insertion of a duplicate copy of a cassette in the conserved array (Escudero et al., 2015). An overview of the mechanisms of integron activity are presented in Figure 1a.

**Figure 1:**
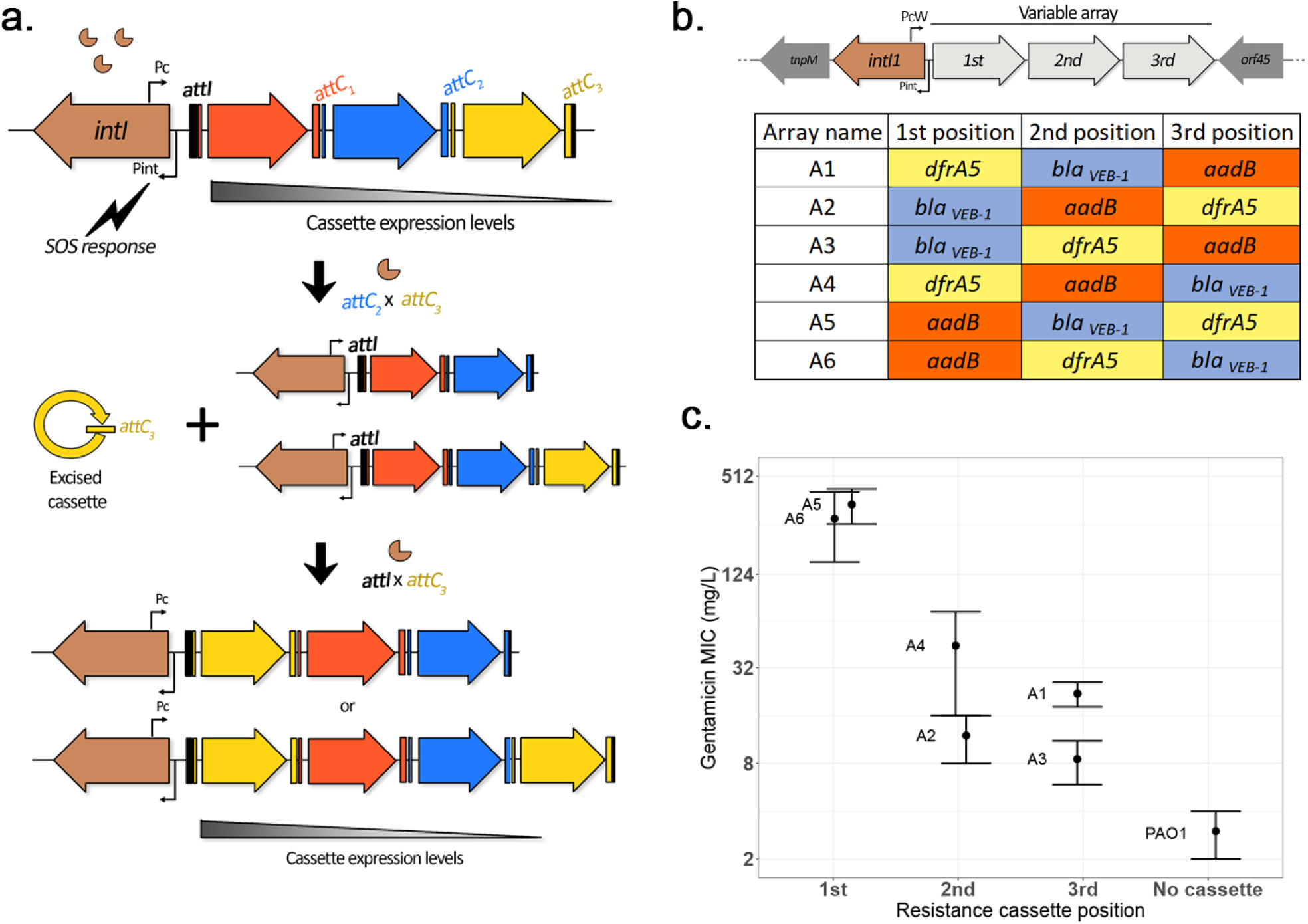
Overview of integron system. a) Diagram of the integron mechanism: the integron consists of an integrase gene, *intI*, followed by an array of promoterless gene cassettes (represented here by arrows). Cassettes are expressed from the Pc promoter within the integrase gene, with decreasing cassette expression along the array. Following the induction of the SOS response, the integrase enzyme promotes cassette excision (recombination between a cassette *attC* site and the *attC* of the preceding cassette, causing excision of the cassette into its circular form). Due to the presence of a replication step in the excision process, a copy of the original array is conserved. Re-integration of the circular cassette can then occur through recombination between the cassette *attC* site and the *attI* site located at the start of either array, leading to an apparent ‘cut-and-paste’ recombination if the cassette integrates in the excised array, or can be assimilated to a ‘copy-and-paste’ reaction if it integrates in a conserved copy of the array. By shuffling and duplicating cassettes the integron has the potential to quickly modulate cassette expression levels. b) Custom integron arrays: the native integron array of the R388 plasmid was replaced by the custom integron arrays WTA1 to WTA6 containing three integron cassettes in every possible order c) Effect of position of the *aadB* cassette on gentamicin resistance levels. Errors bars represent standard error (n=2 to 4)

Due to the stress-inducible regulation of integrase activity, integrons have been proposed to accelerate bacterial evolution by providing ‘adaptation on demand’ (Escudero et al., 2015). According to this hypothesis, integrase-mediated cassette reshuffling in stressful environments allows bacteria to optimize cassette expression and maximize fitness: useful cassettes can be brought forward to ensure maximal expression, while unnecessary cassettes can be kept at the end of the array as a low cost memory of once-adaptive functions, ready to be brought forward when needed (Escudero et al., 2015). Stress-inducible regulation also helps to minimize the costs associated with integrase expression (Lacotte et al., 2017; Starikova et al., 2012), which are thought to come from increased genomic instability created by off-target integrase activity (Harms et al., 2013). Although antibiotics have diverse modes of action, many of the most common classes of antibiotic cause DNA damage that induces the SOS response (Kohanski et al., 2010). This link between antibiotic exposure and integrase activity suggests that cassette re-reshuffling may allow pathogenic bacteria to rapidly adapt to novel antibiotic challenges.

While the molecular mechanisms of integron shuffling are known in detail, our ability to understand the evolutionary benefits provided by this fascinating genetic platform is limited by our understanding of the population biology of integron-mediated antibiotic resistance. For example, constitutive over-expression of the integrase enzyme has been shown to accelerate the evolution of chloramphenicol resistance through the loss of cassettes between Pc and the resistance cassette as well as the formation of co-integrates between integron copies (Barraud & Ploy, 2015). However, to the best of our knowledge, the benefits of cassette shuffling under the integrase natural promoter and its associated LexA regulation have never been investigated. This is an important limitation, as parameters such the re-insertion rate of excised cassettes and fitness costs of integrase expression are predicted to have a strong impact on the evolutionary benefits of the integrase (Engelstädter et al., 2016). Moreoer, integron cassette shuffling has rarely been studied in the large, natural plasmids where class 1 integrons often occur.

Here we directly test the ‘adaptation on demand’ hypothesis using experimental evolution in populations of *Pseudomonas aeruginosa* carrying a variant of the broad host range plasmid R388. We replaced the naturally occurring class 1 mobile integron in R388 with a customized integron containing 3 antibiotic resistance cassettes: *dfrA5* (a trimethoprim resistant dihydrofolate reductase (Sundström et al., 1988)), *bla*_*VEB-1*_ (an extended-spectrum-lactamase (Poirel et al., 1999)) and *aadB* (an aminoglycoside-2’-adenylyltransferase (Cameron et al., 1986)). To directly investigate the role of integrase activity in this system, we also generated a truncated integrase mutant that allows normal levels of cassette expression, but not recombination. Using this system, we found that integrase activity leads to rapid and extensive cassette reshuffling in response to strong selection for increased gentamicin resistance. Specifically, integrase activity caused the insertion of duplicate copies of *aadB* cassettes in the first position of the integron, followed by the loss of redundant cassettes. Crucially, this accelerated the ability of populations to adapt to antibiotic stress, providing good support for the ‘adaptation on demand’ hypothesis. Finally, we show that rapid duplications can also occur with *bla*_*VIM-1*_ cassettes, which confer resistance to ‘last line of defense’ carbapenem antibiotics, in a recently isolated clinical plasmid under meropenem selection.

## Results

### A combinatorial, three-cassette integron system to investigate the impact of cassette position

We replaced the naturally occurring class 1 integron of plasmid R388 with all 6 possible configurations of a class 1 integron containing 3 antibiotic resistance cassettes, including *dfrA5, aadB* and *bla*_*VEB-1*_, and transformed our integron variants into *P. aeruginosa* PA01 (Figure 1B). Integrons have played an important role in the evolution of antibiotic resistance in the opportunistic pathogen *P. aeruginosa*, and are highly prevalent in *P*. *aeruginosa* high-risk clones (Oliver et al., 2015).

As expected, resistance levels declined as cassettes were moved further away from the integrase, and this effect was particularly strong for the *aadB* cassette, which confers resistance to gentamicin (Figure 1C). Interestingly, the relationship between *aadB* position and resistance was not linear: we observed a 6- to 24-fold difference in MIC between arrays containing *aadB* in first and second position, but a less than two-fold difference between arrays with *aadB* in second and third place. In order to investigate the mechanisms behind this trend, we measured the *aadB* cassette transcription levels of the different arrays. Instead of a sharp drop, we observed a steady decrease depending on the cassette position (Figure S1a). Previous work has shown that two short open reading frames contained within the *attI* site can substantially enhance the translation of a cassette lacking a Shine-Dalgarno (SD) sequence when the cassette is located in first position (Hanau-Berçot et al., 2002; Papagiannitsis et al., 2017), showing that cassette position can also modulate translation levels (Hanau-Berçot et al., 2002; Jacquier et al., 2009). Interestingly, the *aadB* cassette contains a reduced SD box (Figure S1b), suggesting that the steep gradient in gentamicin resistance between first and second position was mostly due to decreased translation.

### Integrase activity accelerates the evolution of antibiotic resistance

Given the strong effect of *aadB* cassette position on gentamicin resistance, we decided to use this combination of cassette and antibiotic to experimentally test the hypothesis that integrase activity accelerates resistance evolution. To properly measure the effect of integrase activity on evolvability, we constructed a Δ*intI1* mutant of the A3 array lacking 818bp of *intI1* (total length is 1014bp) but conserving the Pc and Pint promoters. We challenged independent populations of WTA3 and Δ*intI1*A3 with increasing doses of gentamicin using an ‘evolutionary ramp’ design (Gifford et al., 2018; San Millan et al., 2016). Importantly, we did not detect any difference in initial gentamicin resistance (MICexp = 24mg/L) between strains with array A3 or the ΔintI1A3 mutant in the conditions of the evolution experiment (see Material and Methods). We passaged 65 populations of each strain, starting at 1/8 MIC (i.e. 3mg/L) and doubling the concentration of gentamicin each day until reaching 1024 times (24.5 g/L) the initial MIC (Figure 2a). As controls, 15 populations of each strain were passaged without antibiotic (no selection for gentamicin resistance) while 15 populations were passaged at a constant dose of 1/8 MIC (3mg/L) to generate weak selection for gentamicin resistance and plasmid maintenance.

**Figure 2:**
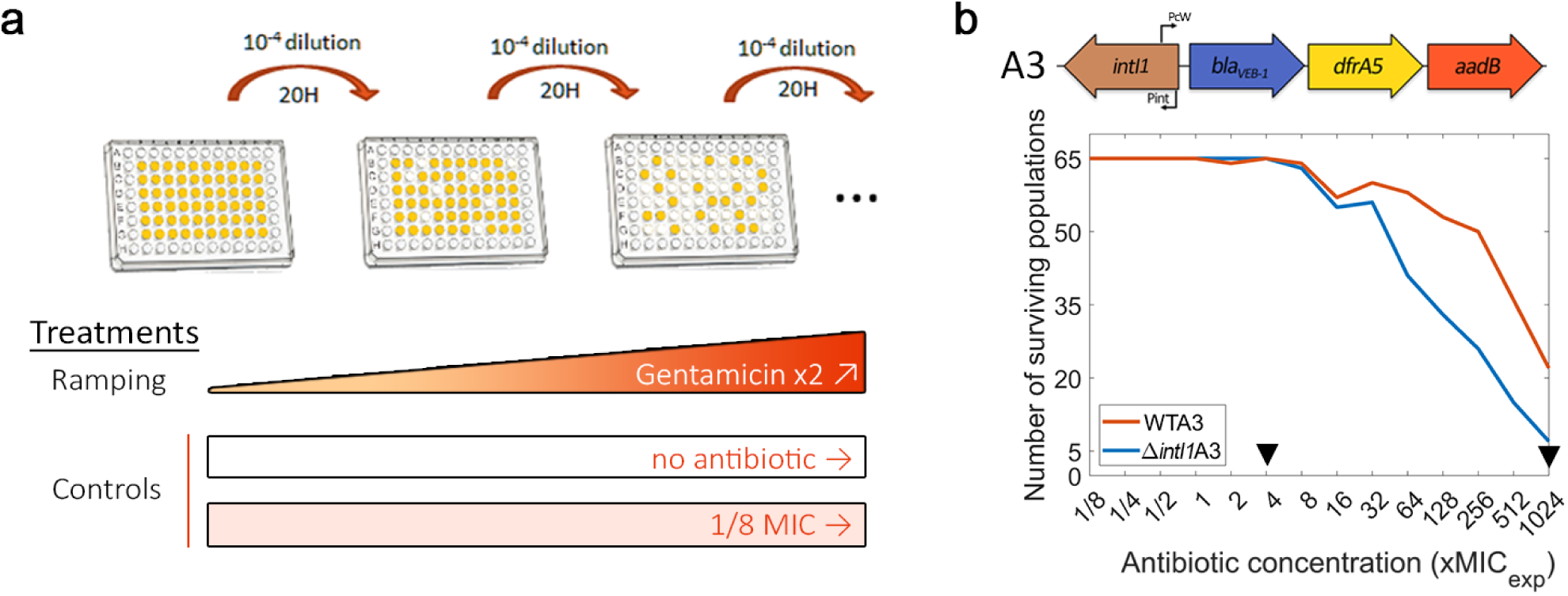
Integrase activity can increase bacteria evolvability against antibiotics. a) Schematic representation of the experimental evolution protocol: b) Top: representation of the WTA3 integron. Bottom: Survival curves of the PAO1:WTA3 and PAO1:Δ*intI1*A3 populations during ramping treatment, monitored using OD595. The black triangles represent time-points where populations were sequenced using whole genome sequencing.

The rapid increase in antibiotic concentration in the ‘evolutionary ramp’ treatment ensures that populations must either evolve increased resistance or face extinction (once concentrations exceed the MIC of the parental strains). Given this, measuring the rate at which populations go extinct provides a way to measure the evolvability of a strain. Crucially, populations of WTA3 populations with a functional integrase had a higher survival rate than those of the Δ*intI1*A3 mutant, showing that the integrase can increase evolvability for antibiotic resistance (Figure 2b; log-rank test: Chisq= 17.7, df=1, p = 3e-05). We did not detect any extinctions in either of the controls, showing that the higher extinction rate ofΔ*intI1*A3 populations was driven by exposure to high doses of gentamicin. To understand the mechanisms by which integrase activity accelerates evolution, we sequenced DNA extracted from randomly chosen populations at a mid-point of the experiment (4XMIC; n=24 WTA3 and 22 Δ*intI1*A3) and all populations that survived until the end of the experiment (1024XMIC; n=21 WTA3 and 6 Δ*intI1*A3 populations).

### Integron evolution under antibiotic treatment

We found evidence for widespread cassette re-arrangement in WTA3 populations and identified 5 novel integron structures that were formed by insertion of the *aadB* cassette and/or deletions of *bla*_*VEB-1*_*-dfrA5* (Figure 3 a,b and Supplementary Table S3 and S4). The junction sites for cassette insertions and deletions were consistent with integrase activity: recombination happened at the 5’-GTT-3’ triplet of the *attI, aadB attC* and *dfrA5 attC* sites. We did not find any evidence for cassette re-arrangements in Δ*intI1*A3 populations, or in control WTA3 populations that we selected at a low dose of gentamicin (1/8X MIC). Cassette re-arrangements were found in most populations at the 4X MIC timepoint, and approximately 90% of populations (19/21) contained cassette re-arrangements by the end of the experiment, highlighting the importance of integrase activity in resistance evolution. Integron structural polymorphisms were found in 50% of populations (12/24) at 4XMIC, but this within-population diversity was transient and almost all populations contained a single dominant integron structure by the final time-point.

**Figure 3:**
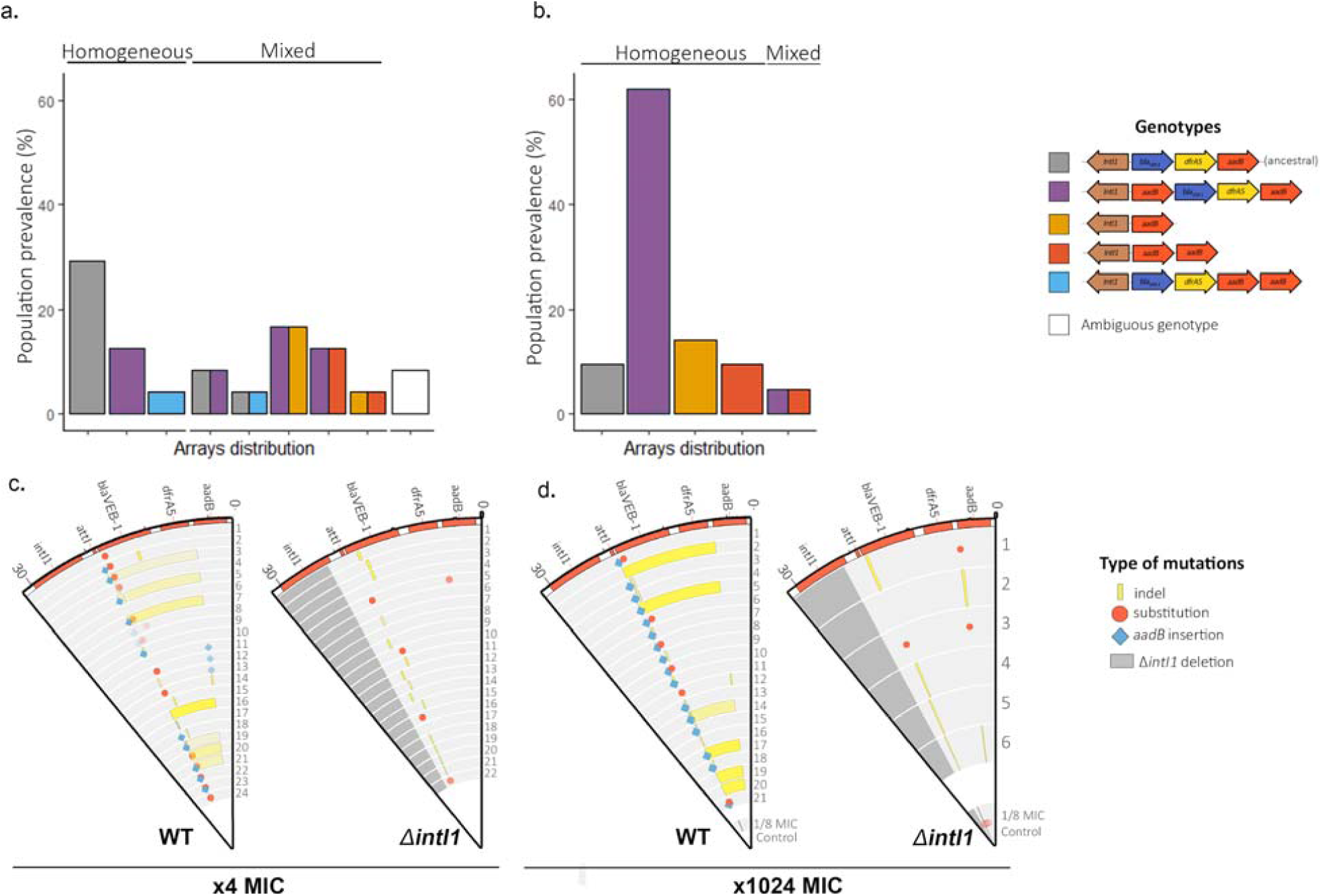
Extensive cassette rearrangements are linked with integrase activity. (a) & (b) Distribution of cassette rearrangements at 4xMIC (a) and 1024xMIC (b) time-points in the WTA3 populations. Homogeneous populations represent populations where only one type of array could be identified while mixed populations contain different arrays as indicated by the corresponding colours. Ambiguous populations correspond to rearrangements that could not be identified with confidence from short-read data. No rearrangement was found in the Δ*intI1*A3 populations. (c) & (d) Representation of the plasmid mutations and rearrangements in the surviving PAO1:WTA3 and PAO1: Δ*intI1*A3 populations at x4 (c) and x1024 MIC (d), mapped to the integron reference sequence. Each circle represents a separate population, with the inner circle representing the variants present in an equimolar pool of six 1/8 MIC control populations. Indels are represented in yellow and single nucleotide substitutions in red. *aadB* insertions are represented by blue lozenges. The color intensity represents the frequency of the corresponding mutation/recombination. The dark grey area in the PAO1:Δ*intI1*A3 populations represents the location of the *intI1* deletion.

The most common novel integron structure contained a ‘copy and paste’ insertion of *aadB* in first position via *attI* × *attC*_*aadB*_ recombination (ie *aadB-bla*_*VEB-1*_*-dfrA5-aadB*). This novel integron should be associated with a large increase (32 fold) in gentamicin resistance due to the dominant effect of first position on *aadB* (Figure 1). Interestingly, we did not identify any *aadB -bla*_*VEB-1*_ *-dfrA5* arrays, which would be the result of an *aadB* excision followed by reintegration of *aadB* in first position within the same array, highlighting the prevalence of ‘copy and paste’ cassette insertions. Degenerate integrons that lack the *bla*_VEB-1_ and *dfrA5* cassettes (i.e. either *aadB* or *aadB-aadB arrays*) were also present at relatively high frequency at both the 4x and 1024x MIC time points. Interestingly, in mixed arrays populations, these reduced arrays were always observed in conjunction with the *aadB – bla*_*VEB-1*_ *- dfrA5 – aadB* array and never with the ancestral array. This repeated association provides good evidence that degenerate arrays evolved via *aadB* insertion in first position, to form the common *aadB – bla*_*VEB-1*_ *- dfrA5 – aadB* array, followed by the *en bloc* deletion of the other cassettes (*bla*_*VEB-1*_ *- dfrA5 – aadB* or *bla*_*VEB-1*_ *- dfrA5*) to form *aadB* and *aadB – aadB* arrays. Recombination happened at the 5’-GTT-3’ triplet of the R box of the *aadB attC* and *dfrA5 attC* sites, suggesting that these deletions were driven by integrase activity, although we cannot rule out the possibility that the *bla*_*VEB-1*_ *- dfrA5 – aadB* cassette deletion was driven by homologous recombination between *aadB* cassettes. The relative prevalence of these two degenerate arrays did not change between the 4X and 1024X time points (4 *aadB* against 3 *aadB-aadB* arrays at 4X MIC, and 3 *aadB* against 3 *aadB-aadB* arrays at X1024 MIC), which suggests that the second *aadB* cassette in the *aadB – aadB* array is redundant. Finally, we found arrays containing a duplicate copy of *aadB* at the end of the array (*bla*_*VEB-1*_ *- dfrA5 – aadB – aadB*), which are likely to have been formed by the insertion of an *aadB* cassette in the middle or at the end of the array through the less frequent *attC* × *attC* integration (intermolecular) reaction. These arrays were only found at the 4X MIC time point, strongly suggesting that they conferred a small increase in gentamicin resistance that was ultimately an evolutionary dead end under strong selection for elevated resistance.

In addition to changes in integron structure, we found widespread integron evolution by mutations in both the WTA3 and Δ*intI1*A3 populations. Mutations in *bla*_VEB-1_ were found in more than 80% of WTA3 and Δ*intI1*A3 populations from the 4x MIC time point, and in almost all populations where the *bla*_*VEB-1*_ cassette was maintained at the 1024x MIC time point, including 5/6 Δ*intI1*A3 and 16/16 WTA3 populations. All mutations in *bla*_*VEB-1*_ were amino acid substitutions (n=9) or indels (n=16) and the 23 amino-acid signal peptide was a hotspot for mutations (12 out of 25 *bla*_*VEB-1*_ mutations), suggesting strong selection to eliminate the secretion of this redundant resistance protein (Supplementary Table S3 and S4). Furthermore, *bla*_VEB-1_ mutations were also found in the ⅛ MIC controls, demonstrating that these mutations were beneficial under low doses of gentamicin, as we would expect if this cassette imposed an important fitness cost. It is unclear if this cost of *bla*_*VEB-1*_ was driven by the presence of gentamicin (i.e. collateral sensitivity), because the entire R388 plasmid was lost in every control population that was passaged in antibiotic-free medium. Parallel evolution also occurred in the intergenic region between the *dfrA5 attC* site and the start codon of the *aadB* cassette. These mutations were very rare at the 4XMIC time points (2/46 populations), but were present at a high frequency in Δ*intI1*A3 populations from the final time point (4/6 populations). We speculate that these mutations were favored in Δ*intI1*A3 populations as they increased the translation rate of the weakly expressed 3^rd^ position *aadB* cassette. Similarly, we identified one 41 bp deletion within the *dfrA5 attC* site of a WTA3 population which may increase a translational coupling with the previous *dfrA5* cassette (as in (Jacquier et al., 2009)) or lead to the creation of a fused protein with part of the previous cassette. Finally, we observed extended deletions in one WTA3 population and in the 1/8 MIC WTA3 pooled control. These deletions occur between *attC*_*aadB*_ and different positions within the plasmid *trwF* gene, effectively deleting most of the genes involved in mating pore formation (Supplementary Figure S3), with the sequence of the junctions sites pointing toward potential off-target activity of the integrase (abundance of 5’- GNT-3’ secondary sites) (Supplementary Figure S4).

### Chromosomal evolution

The integron integrase is known to have off-target effects, suggesting that integrase activity may also have an important effect on chromosomal evolution, for example by recombination between chromosomal *attC* sites or by increasing the bacteria mutation rate.

Chromosomal evolution occurred more rapidly in the Δ*intI1*A3 populations than in the WTA3 populations, as demonstrated by the high cumulative frequency of mutations in Δ*intI1*A3 populations (mean = 1.01; s.e = 0.17, see Supplementary Figure S6) at 4XMIC compared to WTA3 populations (mean = 0.48; s.e = 0.11; t =−2.69, df = 35.83, p = 0.01, Welch Two Sample t-test)). However, accelerated chromosomal evolution in the absence of integrase activity was short-lived, and cumulative frequency of mutations in the WTA3 and Δ*intI1*A3 populations was almost identical at the end of the experiment (mean WTA3 = 2.45 SNPs; mean Δ*intI1*A3 = 2.65 SNPs, see Supplementary Figure S5). Crucially, we found only one case of chromosomal recombination, with a 3.5 kb deletion between the two highly homologous ccoN1 and ccoN2 cytochrome C subunits in one WTA3 population (Supplementary Table S3), showing that off-target effects of the integrase were undetectable in our experiment, in spite of our extensive genomic sequencing.

In total, we identified 41 different SNPs and 58 short indels in 8 intergenic regions and 41 genes, with a similar spectrum of mutations in the ramping WTA3 and Δ*intI1*A3 populations (Figure 4; Supplementary Table S3 and S4). Several lines of evidence indicate that the overwhelming majority of mutations were beneficial mutations that reached high frequency as a result of selection. First, many of the mutated genes have known roles in antibiotic resistance; for example, 11/41 mutated genes have also been identified in an aminoglycoside resistance screen in *P. aeruginosa* (Schurek et al., 2008). Second, parallel evolution was very common. Repeated evolution occurred in 11 of 42 (26%) genes and 3 of 8 (38%) intergenic regions and 68% of mutations occurred in these genes. Only 1 out of 41 mutations in coding regions was synonymous, providing evidence that the rapid evolution of proteins was driven by positive selection, and not simply by an elevated mutation rate. Finally, we found almost no overlap between the genes that were mutated in the ramping populations and the controls, implying that the evolutionary response of the ramping populations was dominated by selection for high levels of gentamicin resistance.

**Figure 4:**
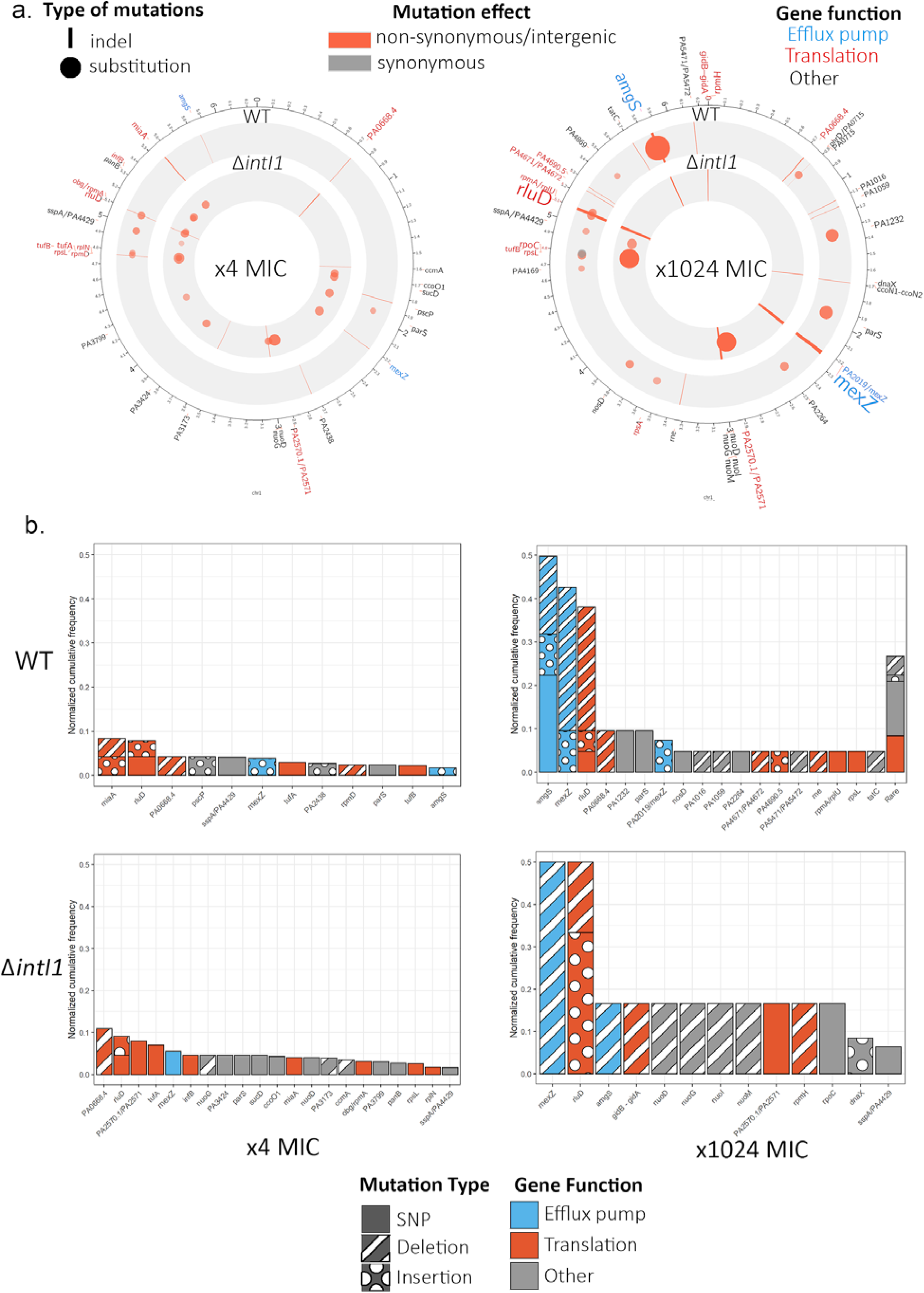
Chromosome evolution. a) Summary of the chromosomal mutations at x4 MIC (left) and at x1024 MIC (right) mapped to the PAO1 reference sequence. Each circle represents a summary of each genotype. The type (indel, substitution) of mutation for each gene is represented by the shape of the marker (line, circle), while the marker colour represents the effect of the mutations (nonsynonymous/ intergenic vs synonymous) and its colour intensity and size represents its normalized cumulative frequency per gene. The size of the gene labels on the outer ring represents the overall cumulative frequency of mutations present in this gene across all populations from this time-point. b) Cumulative frequency of mutations for each gene normalized by the number of populations within each genotype and time-point. Genes are colored by resistance mechanism and the type of the mutations (single nucleotide substitution, insertion, deletion)) is indicated by the patterning.

The initial stages of adaptation to gentamicin were driven by mutations in a very diverse set of genes, with a strong bias towards genes that are involved in translation, such as *rluD* and PA0668.4, which encodes for the 23S ribosomal RNA (Figure 4). Interestingly, we observed divergent mutational trajectories of evolution in the WTA3 and Δ*intI1*A3 backgrounds: the number of genes that were mutated in both backgrounds was small (n=7) relative to the total number of mutated genes in either background (n=26) and the correlation in mutation frequencies across backgrounds was weak (rho=0.029; P=0.89 – Spearman test).

Continued selection for elevated gentamicin resistance resulted in two changes in chromosomal evolution (Figure 4). First, chromosomal evolution became dominated by mutations in a few key target genes, implying that many of the trajectories of chromosomal evolution followed during early adaptation ultimately led to evolutionary dead ends. For example, the correlation in mutation frequencies between early and late time points was very weak, in both WTA3 (rho =−0.10, P=0.58) and Δ*intI1*A3 (rho=0.20, P=0.307). In particular, we found evidence of extensive parallel evolution in *mexZ, amgS*, and *rluD* in both the WTA3 and Δ*intI1*A3 populations. At a functional level, the mutations found at 1024XMIC are predominantly involved in antibiotic efflux, as opposed to translation. *mexZ* is a transcription factor that represses the expression of the *mexXY* multidrug efflux pump operon. Mutations inactivating *mexZ* cause a 2 to 16 fold increase in aminoglycoside resistance, and have been widely identified in aminoglycoside resistant *P. aeruginosa* isolates found in cystic fibrosis patients (Vogne et al., 2004). *AmgS* is part of an envelope stress-responsive two-component system *AmgRS* (Lau et al., 2013), and *amgS* mutations upregulate the *mexXY* multidrug efflux system in the presence of aminoglycosides (Lau et al., 2015).

### Cassettes duplication in a clinically relevant plasmid

Resistance to carbapenem antibiotics in *P. aeruginosa* has emerged as an important clinical threat; for example, the WHO has designated carbapenem resistant *P. aeruginosa* as a ‘critical priority’ for the development of new antibiotics. Interestingly, mobile integrons carrying multiple bla_VIM-1_ carbapenemase cassettes have been found in clinical isolates of *P. aeruginosa* (San Millan, Toll-Riera, et al., 2015), suggesting that cassette duplications may play an important role in clinical settings. To test this idea, we challenged 30 populations of *P. aeruginosa* carrying a plasmid (pAMBL-1), which contains an integron carrying a single copy of bla_VIM-1_ followed by *aadB*, with increasing doses of meropenem using a similar evolutionary ramp experiment (Supplementary Figure S7). PCR screening of populations that survived at x2 MIC identified numerous cassette rearrangements of both the *bla*_*VIM-1*_ and *aadB* cassettes, with potential *bla*_*VIM-*_ *1* duplications occurring in all 14 surviving populations (Supplementary Figure S7). Short-read sequencing of clones isolated from two of these populations confirmed the presence of duplications, as demonstrated by increased copy number of bla_VIM-1_ per plasmid (2.0 copies/plasmid (95% CI: 1.90 – 2.10); and 2.67 copies/plasmid (95% CI: 2.53 – 2.78)). Although it is not possible to definitely prove the role of the integrase without control populations lacking a functional integrase, these results strongly support the idea that ‘copy and paste’ cassette re-arrangements can drive the rapid evolution of elevated carbapenem resistance and was the mechanism behind the bla_VIM-1_ amplification observed in the plasmid pAMBL2 isolated in the same hospital (San Millan, Toll-Riera, et al., 2015).

## Conclusion

Mobile integrons are widespread genetic platforms involved in the interchange and expression of antibiotic resistance cassettes in bacteria. Antibiotic-induced cassette re-shuffling mediated by the SOS response (Barraud & Ploy, 2015; Cambray et al., 2011; Guerin et al., 2009) has been proposed to increase bacterial evolvability by providing ‘adaptation on demand’ to newly encountered antibiotics (Engelstädter et al., 2016; Escudero et al., 2015). We tested this hypothesis by quantifying the impact of integrase activity on adaptation to increasing doses of gentamicin in populations of *P. aeruginosa* carrying a customized integron on a broad host range plasmid. Crucially, integrase activity accelerated the evolution of gentamicin resistance through rapid and repeated re-shuffling of the *aadB* resistance cassette, providing experimental support for the ‘adaptation on demand’ hypothesis.

We observed a diversity of cassettes rearrangements as a result of integrase activity, whose diverse prevalences can help us understand the evolutionary dynamics underlying cassette shuffling. Cassette shuffling and duplication were extremely frequent, both with the *aadB* and the *bla*_*VIM-1*_ cassette. The semi-conservative nature of cassette excision (Escudero et al., 2015) implies that cassette re-shuffling can be assimilated to either a ‘cut and paste’ or ‘copy and paste’ process. Strikingly, all of the *aadB* re-arrangements that we observed were the product of ‘copy and paste’ re-shuffling, resulting in the duplication of *aadB* (Supplementary Figure S8). A bias towards evolution by ‘copy and paste’ is expected if increased copy number of the cassette under selection is beneficial. In this case, the benefit provided by *aadB* is strongly dependent on position, suggesting that ‘copy and paste’ insertions are unlikely to have provided stronger benefits than ‘cut and paste’ rearrangements in first position. Furthermore, we did not detect any advantage of *aadB-aadB* as compared to *aadB* arrays, suggesting that secondary copies of *aadB* were redundant in arrays containing an *aadB* cassette in first position. The presence of multiple integrons within the same cell may also create an apparent bias towards ‘copy and paste’ re-shuffling: if re-insertion of an excised cassette is equally likely between all the integron copies present in a cell, we would expect the chance of an excised *aadB* cassette to re-insert into its original array to be 25% to 30% given the copy number of the R388 plasmid (2-3 per cell (Fernández-López et al., 2006)). However, the absence of any ‘cut and paste’ reaction product in our experiments, instead of the expected 30%, suggests that the integrase enzyme may be inherently biased towards ‘copy and paste’ cassette re-arrangement, potentially by favoring the reinsertion of an excised cassette into the conserved array.

Although duplicated cassettes are relatively common in large chromosomal integrons, such as the *V. cholerae* super integron (Escudero et al., 2015), they are rarely found in mobile integrons (San Millan, Toll-Riera, et al., 2015; Stokes & Hall, 1992). For example, duplicate cassettes are only found in 5% of the integrons that contain 2 or more cassettes in the INTEGRALL database (Moura et al., 2009) (Supplementary Table S5). Difficulties associated with resolving duplications from short-read sequencing data probably contribute to this (Alkan et al., 2011), but it is clear that duplicate cassettes are rare. Interestingly, the duplicate copies of selected cassettes created by ‘copy and paste’ re-shuffling facilitate the loss of redundant cassettes. First, duplication of cassettes with highly recombinogenic *attC* sites, such as *aadB*, facilitates the integrase-mediated excision of redundant cassettes. In this case, the insertion of *aadB* in first position created the opportunity for the loss of the *bla*_VEB-1_-*dfrA5* cassettes, through an *attC*_aadB_ × *attC*_*dfrA5*_ reaction, or the excision of *bla*_*VEB-1*_*-dfrA5-aadB*, through an *attC*_*aadB*_ × *attC*_*aadB*_ recombination. Homologous recombination between duplicate cassettes or between copies of integrons on different plasmids provides a second mechanism for integron degeneration (Andersson & Hughes, 2009), in this case resulting in the formation of a single copy of the duplicate gene. This also highlights how extremely mobile cassettes, such as *aadB*, can compensate for the lack of mobility of other less recombinogenic cassettes, like *bla*_*VEB-1*_ (Aubert et al., 2012). The constitutive expression of cassettes from the Pc promoter ensures that redundant cassettes impose a fitness cost (Lacotte et al., 2017), implying that selection for cassette loss is likely to be common. Given this, we argue that semi-conservative nature of cassette excision is key to the evolutionary benefits of integrase activity, as it allows integrons to rapidly gain additional copies of selected cassettes and facilitates the subsequent elimination of costly redundant cassettes.

In our experiments, mutations in the chromosome and in the integron made an important contribution to resistance evolution and are key to understanding the effective integrase evolutionary benefits. For example, integrase activity was associated with the loss of the redundant *bla*_*VEB-1*_ cassette, which is in line with the integrase-mediated loss of redundant cassettes during selection for elevated chloramphenicol resistance observed by (Barraud & Ploy, 2015). However, the *bla*_*VEB-1*_ cassette was also rapidly inactivated by mutations in Δ*intI1* populations. While integrase activity offers additional evolutionary pathways to alleviate the cost of *bla*_*VEB-1*_, the presence of numerous mutational targets achieving the same effect may offset the observable evolutionary benefit of the integrase. Similarly, mutations in the promoter of the *aadB* cassette provided an alternative mechanism to increase the expression of this cassette that did not depend on integrase activity. Moreover, *P. aeruginosa* has a strong potential to adapt to aminoglycosides via chromosomal mutations (López-Causapé et al., 2018; Schurek et al., 2008). Given our populations possess a sizable evolvability potential through mutations, even in the absence of integrase activity, we hypothesize an even stronger impact of the integrase should be observable in environments or species where evolution possibilities through mutations are more limited. Finally, we detected an interesting interplay between chromosomal evolution and integrase activity, with a divergence in the evolution of the two genotypes at the early x4 MIC time-point, potentially the most clinically relevant. Rapid chromosomal and plasmid evolution via mutations across a wide range of target genes allowed populations lacking a functional integrase to evolve resistance to >MIC concentrations of gentamicin. The wild-type populations, on the other hand, showed a much-reduced mutation prevalence, as chromosomal mutations were potentially out-competed by the frequent and more efficient cassette rearrangements. This accelerated chromosomal evolution in Δ*intI1A3* populations was short-lived, confirming that rapid evolution was driven by more effective selection for mutations in populations lacking an integrase, rather than a difference in mutation rate per se. All populations that were able to survive very high levels of gentamicin exposure evolved by mutations of a common suite of target genes involved in antibiotic efflux and ribosomal modification, highlighting the fact that integrase activity did not ultimately alter the mutational routes to high level resistance, but can impact chromosomal evolution during the early evolution of resistance.

Systems that upregulate the mutation rate under stress are widespread in bacteria (Foster, 2011; MacLean et al., 2013), and beneficial mutations generated by these systems can accelerate adaptation to stress (but see (Torres-Barceló et al., 2015)). However, most of the mutations generated by these systems will be deleterious, and stress-induced mutagenesis will therefore tend to reduce fitness, particularly if the deleterious effects of mutations are exacerbated by stress (Kishony & Leibler, 2003). The integron integrase is known to have off-target activity (Recchia et al., 1994), and it has been argued that the costs associated with off-target recombination contribute to the cost of integrase expression (Harms et al., 2013), limiting the evolutionary benefit of this system (Engelstädter et al., 2016). Although we found one case of a deletion in the R388 plasmid that suggests off-target activity of the integrase, we found no evidence of chromosomal rearrangements or increased mutation rates that could be linked to integrase activity. On the opposite, integrase activity reduced the prevalence of mutations at the 4xMIC time-point, as cassette rearrangements out-competed chromosomal mutations. Unlike stress-induced mutagenesis, which increases mutations across the genome, integrase activity creates variation in a focused manner by creating high levels of variation exclusively in a region of the genome containing genes involved in response to stress, allowing bacteria to benefit from increased diversity without compromising genomic integrity.

In conclusion, our study supports the view that integrons provide bacteria with an incredible opportunity to evolve in response to new antibiotic challenges by rapidly optimizing the expression of cassettes. Integrase activity allows bacteria to rapidly gain additional copies of selected cassettes and eliminate redundant cassettes, while the high specificity of integrase-mediated recombination maintains genomic integrity, minimizing the costs of integrase activity. Given the importance of cassette re-shuffling, we argue that integrase activity will accelerate resistance evolution most for highly mobile cassettes that display strong positional effects, such as *aadB*. Given this, we argue that cassette re-shuffling will be most important in cases where bacteria have limited ability to adapt to antibiotics, for example when only a small number of mutations can increase resistance, or where resistance mutations carry large fitness costs. Integrase activity also provides bacteria with the opportunity to capture new resistance cassettes, an important challenge for future work will be to study the evolutionary processes driving cassette acquisition. Our work also supports the view that treatment strategies should seek to target integrons, for example by combining antibiotics with adjuvants that limit integrase activity by inhibiting the SOS response (Hocquet et al., 2012), or by using combinations of antibiotics that impose conflicting selective pressures on the integron.

## Material & Methods

### Bacteria and growth conditions

A complete list of strains and plasmids can be found in Table S1. Unless stated, bacteria cultures were grown overnight at 37°C with shaking in LB Miller broth (Sigma Aldrich) and supplemented with 100 mg/L of ceftazidime when required to select for the integron-bearing plasmids.

### Strains construction

Six integron arrays covering all possible cassette orders were created using the plasmid R388 (Avila & de la Cruz, 1988) as backbone and three resistance cassettes: *aadB, bla*_*VEB1*_ and *dfrA5*. The *bla*_*VEB1*_ and *aadB* cassettes were amplified from the integron of the *E. coli* MG-1 clinical isolate (Poirel et al., 1999), while the *dfrA5* cassette was obtained from an enteroinvasive *E. coli* strain isolated in Senegal (Gassama et al., 2004). These cassettes were then assembled into custom integron arrays using Gibson assembly and inserted into the plasmid R388 while replacing its original *dhfr – orf9 – qacE*Δ*1 – sul1* integron array (Fernández-López et al., 2006). The original R388 strong PcS promoter variant (high cassette expression but low integrase activity (Jové et al., 2010)) was replaced by the weaker PcW promoter to guarantee high integrase activity. Δ*intI1* mutants of these custom integrons were created by introducing a 948 bp deletion of the integrase *IntI1* gene during array construction, deleting most of the integrase open reading frame but conserving both the Pint and Pc promoters. The final arrays were then first transformed into chemically competent E. coli MG1655.

These plasmids were then conjugated into *P. aeruginosa* through filter mating using the previous *E. coli* strains as donors and PAO1 as recipient. Bacteria were incubated overnight in Luria-Bertani (LB) broth with 100mg/L of carbenicillin at 37°C for the donors and in LB Miller broth without antibiotic at 42°C for the recipient bacteria. The next day cells were spun down, washed, and re-suspended in LB broth, before mixing in a 1:4 donor to acceptor ratio. The mix, as well as pure donor and acceptor controls, were put on filters placed on LB agar without antibiotics and incubated at 37°C overnight. Afterwards, filters were placed in tubes containing LB media and agitated. The resulting supernatants were plated on LB agar supplemented with 50 mg/L of kanamycin and 25 mg/L of ceftazidime and incubated for 48h. As *P. aeruginosa* PA01 has a higher innate resistance to kanamycin than *E. coli* MG1655 due to a chromosomally encoded phosphotransferase (Okii et al., 1983), kanamycin was used to used to select against the *E. coli* donors, while ceftazidime was used to select for the plasmid in the *P. aeruginosa* transconjugants. The final colonies were controlled by PCR for the presence of the plasmid and the absence of *E. coli* DNA.

### Minimum inhibitory concentration (MIC) determination

The minimum inhibitory concentrations (MIC) for each antibiotic were determined in cation-adjusted Mueller-Hinton Broth 2 (MH2), following the broth microdilution method from the Clinical and Laboratory Standards Institute guidelines (CLSI, 2017). Briefly 5x c.f.u bacteria inocula were prepared using individual colonies grown on selective agar in interlaced 2-fold-increasing concentrations of antibiotics and incubated for 20h in a shaking incubator at 37°C.

The next day, plates’ optical density (OD_595)_ was read using a Biotek Synergy 2 plate reader. Wells were considered empty when the overall was under 0.1 and the MIC for each assay was defined as the minimal concentration in which growth was inhibited in all three technical replicates (separate wells, but grown on the same day from the same inoculum). The final MICs values are the average of two to four replicate assays (from separately prepared inocula, on different days).

### Experimental evolution with custom arrays

As antibiotics’ MICs vary depending on the size of the starting inoculum (Brook, 1989), we first determined minimum inhibitory concentrations in densities similar to the experimental evolution experiment (further called MIC_exp_). Overnight cultures inoculated from 2-3 morphologically similar colonies grown on selective agar were incubated for 20h with shaking in MH2 media with antibiotics. These overnight cultures were then diluted 1/10 000 and supplemented with doubling concentrations of gentamicin in three replicates. MIC_exp_ were determined the next day after 20h of incubation using OD_595_ measurements. This process was repeated twice. In these conditions the MIC_exp_ for PAO1:WTA3 and PAO1:Δ*intI1*A3 were identical at 24 mg/L.

At the start of the experiment 90 individual colonies grown on selective agar of each strain (PAO1:WTA3 and PAO1:Δ*intI1*A3) were inoculated in 200 μL of MH2 media supplemented with gentamicin at a concentration of 1/8 MIC_exp_. WT and Δ*intI1* strains were placed in a chequerboard pattern by interlacing the different genotypes to control easily for cross-contamination. Wells at the edge of every plate were kept bacteria-free to avoid edge effects and identify contaminations. These 90 populations were passaged every day with a 1/10 000 dilution factor and the antibiotic concentration was doubled, starting at 1/8 MIC_exp_ until reaching a concentration of 1024x MIC_exp_. Alongside these 90 populations per strain which were transferred in increasing antibiotic concentrations, 30 populations per strain were transferred as controls in constant conditions: 15 without antibiotic and 15 at a constant concentration of 1/8 MIC_exp_. Each population’s OD_595_ was measured each day and a population was considered extinct when its OD_595_ fell below 0.1 after 20h of incubation. All populations were frozen in 15% glycerol every two days.

### DNA extraction

Liquid cultures were grown from the frozen stock of all surviving PAO1:WTA3 and PAO1:Δ*intI1* A3 populations at x1024 MIC_exp_ in LB Miller media supplemented with gentamicin at x128 MIC_exp_, and were incubated for 24h with shaking. Six populations were inoculated from each control treatment in either LB Miller supplemented with a concentration of MIC_exp_ or with no antibiotic. For the x4 MIC_exp_ time point 26 populations of each PAO1:WTA3 and PAO1:ΔA3 genotype were regrown in x2 MIC_exp_ concentration of gentamicin. Ancestral PAO1:WTA3 and PAO1:Δ*intI1*A3 populations were incubated in 100 mg/L of ceftazidime from the initial frozen stock. DNA extractions of the whole populations were carried out using the DNeasy Blood & Tissue Kit (Qiagen) on the QiaCube extraction platform (Qiagen) combined with RNAse treatment. DNA concentrations were determined using the Quantifluor dsDNA system (Promega).

### PCR controls and analysis

At the x1024 and x4 MIC_exp_ transfers all surviving populations were controlled for cross-contamination by verifying the size of the integrase by PCR (Primers given in Table 2). Starting materials were either 2 μL of extracted DNA (x1024 MIC_exp_ time point) or 2 μL of inoculate previously incubated for 24h then boiled for 10 minutes at 105°C (x4 MIC_exp_ time point). PCR reactions were carried out using the GoTaq G2 DNA mastermix (Promega) and the following protocol: 30s at 95°C, 30s at 55°C, 3 minutes at 72°C for 30 cycles. Plate mishandling during the transfers resulted in the contamination of 40 and 34 wells out of 180 ramping populations for each array. Areas of the plates where cross-contamination was detected in several wells in close proximity were excluded from the rest of the analysis, for a final population number of 65 per strain. The final populations at x1024 MIC_exp_ were analysed by PCR to determine the position of the *aadB* cassette relative to the start and the end of the array as well as identify any *aadB* duplications or inversions and deletions of the plasmid backbone.

### Next Generation Sequencing and bioinformatic pipeline

Library preparation and Next-Generation sequencing using the NovaSeq 6000 Sequencing System (Illumina) was carried out at the Oxford Genomics Centre at the Wellcome Centre for Human Genetics. 22 PAO1:WTA3 and 6 PAO1:Δ*intI1*A3 populations were sequenced from the x1024 MIC_exp_ time point. For each control treatment, six populations were pooled together and sequenced as one. For the x4 MIC_exp_ time point 26 PAO1:WTA3 and 26 PAO1:Δ*intI1*A3 were sequenced.

PCR duplications and optical artifacts were removed using MarkDuplicates (*Picard Toolkit*, n.d.) then low quality bases and adaptors were trimmed from the sequencing reads using Trimmomatic v0.39 (Bolger et al., 2014). Finally overall read quality control was performed using FastQC (Simon Andrews, 2010) and multiQC (Ewels et al., 2016). During this process one PAO1:WTA3 sample from the x1024 MIC time-point was removed due to the presence of non-*Pseudomonas* DNA.

Single-nucleotide polymorphisms (SNP), point insertion and deletion identification was performed using the breseq (Barrick et al., 2014) pipeline in polymorphism mode. For each population reads were first mapped to the *P. aeruginosa* PAO1 complete genome NC_ 002516.2 and the predicted sequence of WTA3. Non-mapped reads from the unevolved PA01:WTA3 population were then assembled *de novo* using SPAdes (Bankevich et al., 2012) and any open-reading frame identified using Prokka (Seemann, 2014) and further used as an additional reference to map unaligned reads from the other populations. The pipeline output was then further processed in MATLAB (MathWorks). Variants present in the un-evolved ancestor populations at any frequency were filtered out. We also excluded variants reaching a frequency of less than 30% within a single population. A 5% threshold was applied to the pooled controls, which allows the detection of any variant present in more than 30% of a single population. Final results were exported in table format and processed for visualisation using Circos (Krzywinski et al., 2009) and Geneious (Biomatters). Apart from the expected *intI1* deletion, the PA01:Δ*intI1*A3 genome was shown to differ from PA01:WTA3 and PA01 by two SNPs likely to have arisen during the conjugation process: one in PA3734 and one in the phzM/phzA1 intergenic region. PA3734 is predicted to be a lipase involved in cell-wall metabolism (Dettman et al., 2015) and may be involved in quorum-sensing (Levesque, 2006), while phzM and phzA1 are involved in the production of pyocyanins (Higgins et al., 2018). No literature linking those genes to aminoglycoside resistance was identified. Four PA01:Δ*intI1*A3 samples and one PA01:WTA3 sample from the 4x MIC time point were removed from the analysis due to a wrong or mixed genotype from potential mislabelling or mishandling during DNA processing, leading to a final genomic dataset of 22 PA01:Δ*intI1*A3 and 24 PA01:WTA3 populations at 4x MIC. All samples from the 1024x MIC time point were of the correct genotype.

Potential new junctions between distant regions of the reference genome were identified through the *breseq* software on the plasmid and on the chromosome (Barrick et al., 2014). Copy number variants were identified with CNOGpro (Brynildsrud, 2018) and used to confirm potential duplications or large scale deletions. Finally, *de novo* assembly of the plasmids using plasmidSPAdes (Antipov et al., 2016) was carried out to provide additional evidence for the cassette rearrangements and visualised using Bandage (Wick et al., 2015). The robustness of the detection of cassette rearrangement from the sequencing data was tested by cross-referencing the predicted recombinations with the results from the PCR screen at x1024 MIC: all predicted recombinations matched the bands of the PCR screen, and only two extraneous bands could not be explained in three populations (Supplementary Figure S2), confirming the robustness of the bioinformatic analysis.

### Statistical analysis

Statistical analysis was carried out using R (version 3.6.1) and RStudio (Version 1.2.5). Survival analysis using the log-rank test was performed using the survival (Therneau, 2020) package to compare survival rates between populations with and without a functional integrase.

## Acknowledgements

We thank Laurent Poirel and Marie-Cecile Ploy for the gift of the strains containing the cassettes used in the construction of the integrons arrays. We are grateful to Natalia Kapel and Gerda Kildisiute for experimental support and Julio Diaz Caballero and Jessica Hedge for bioinformatic support. This project was funded by Wellcome Trust Grant 106918/Z/15/Z held by R.C.M. We thank the Oxford Genomics Centre at the Wellcome Centre for Human Genetics (funded by Wellcome Trust grant reference 203141/Z/16/Z) for the generation and initial processing of the sequencing data. C.S. was supported by funding from the Biotechnology and Biological Sciences Research Council (BBSRC) [grant number BB/M011224/1]. J.A.E. was supported by the European Research Council (ERC) through a Starting Grant (803375), the *Atracción de Talento* Program of the *Comunidad de Madrid* (2016-T1/BIO-1105) and the *Ministerio de Ciencia, Innovación y Universidades* (BIO2017-85056-P)

## Author contributions

J.A.E. and R.C.M. conceived the project and J.A.E, C.S. and R.C.M conceived the experimental design. C.S. and J.A.E performed and analyzed the experiments. C.S. performed the bioinformatic analyses. C.S., J.A.E and R.C.M wrote the paper.

## Competing financial interests

The authors declare no competing financial interests.

## Data availability

The sequences generated in this work have been deposited in the European Nucleotide Archive database under the accession code [PRJEB40301]. All other datasets generated during this study are available from the data repository Dryad under the doi:10.5061/dryad.rv15dv469.

## Supplementary Information

### Supplementary Materials

#### aadB cassette transcription levels

##### RNA & DNA extractions

Each bacterial strain was inoculated in MH2 medium supplemented with antibiotics and grown overnight at 37°C with constant shaking (225 rpm). The overnight cultures were diluted 1:50 in fresh MH2 without antibiotics and incubated until they reached an OD600 between 0.5 and 0.6. Both RNA and DNA were extracted for each sample. Half of each culture was mixed with RNAprotect Bacteria Reagent (Qiagen) according to the manufacturer instruction. Total RNA extraction was performed using the RNeasy Mini kit (Qiagen) on the QIAcube extraction machine (Qiagen). The other halves were treated with RNase and used to extract total gDNA using the DNeasy Blood & Tissue Kit on the QiaCube (Qiagen). Each strain was extracted three times from cultures started on different days.

##### Plasmid copy number

Plasmid copy number was determined for each gDNA sample using the approach described in (San Millan, Santos-Lopez, et al., 2015): all samples were first digested in order to linearize the plasmid using the restriction enzyme BamHI (BamHI FastDigest, Thermo Fisher Scientific) according to the manufacturer instruction for gDNA digestion. Linearizing the plasmid increases DNA template accessibility and therefore prevents from underestimating the plasmid copy number (Providenti et al., 2006). The amplified regions were controlled for the absence of BamHI restriction sites. The orf9 gene was used as the R388 plasmid target and the mono-copy rpoD gene was used as chromosomal target for *P. aeruginosa* (Primers given in Supplementary Table S2). Amplifications were carried out using the Luna Universal Probe qPCR Master Mix (New England Biolabs). Thermal cycling protocol consisted of 1s at 95°C (Denaturation), 20s at 60°C (Annealing/Extension) for 40 cycles. Melting curve analysis was included for samples detected without probes. 4-fold dilution standard curves were included to control for differences in primer efficiencies. Plasmid copy number was calculated as the ratio between the plasmid and chromosomal target DNA quantities.

##### Reverse transcription and qPCR

All RNA samples were treated with the TURBO DNA-free Kit (ThermoFisher) to eliminate genomic DNA. Concentration of the RNA samples was quantified using the Quantifluor RNA system (Promega). cDNA was synthesized from 100ng of RNA templates using random primers from the GoScript Reverse Transcription Mix (Promega). qPCR was carried out on the StepOnePlus Real-time PCR platform (Applied Biosystems) using the iTaq Universal SYBR Green Supermix. The cassette, as well as two reference genes (actpA and acp), were amplified using the primers described in Supplementary Table S2 in two technical replicates for each extraction. Standard curves for the pair of cassette primers was included in each PCR using restriction enzyme-digested gDNA as template, and used to quantify the amount of target cDNA in each sample to control for inter-run variations. Melting curve analysis was included after each run to test for non-specific amplification products. For each biological replicate the cassette transcript levels were normalized based on the geometric means of the two internal reference genes, using the first array A1 as a reference, before division by its plasmid copy number.

#### Experimental evolution with pAMBL1

Thirty colonies were inoculated in 100 μL of MH2 broth and transferred every day in doubling concentrations of meropenem with a 1/10 000 dilution and frozen in 15% glycerol every other day. Population survival was monitored each day after reaching a concentration of x1 MIC_exp_ by plating every well on a MH2 agar plate without antibiotic using a pin replicator. Extinction of a population was defined as the absence of a visible colony after 24h incubation at 37°C.

Surviving populations at x2 MIC_exp_ were grown on a MH2 agar plate without antibiotic and used as substrate for PCR. Primers were used to identify potential duplications of the bla_VIM_ cassette by PCR. PCR reactions were carried out using the GoTaq G2 DNA mastermix (Promega) and the following protocol: 30s at 95°C, 30s at 55°C, 3 minutes at 72°C for 30 cycles. Single clones from two different populations were sequenced through whole genome sequencing and analysed using the same protocol as described previously.

## Supplementary figures

**Fig S1.**
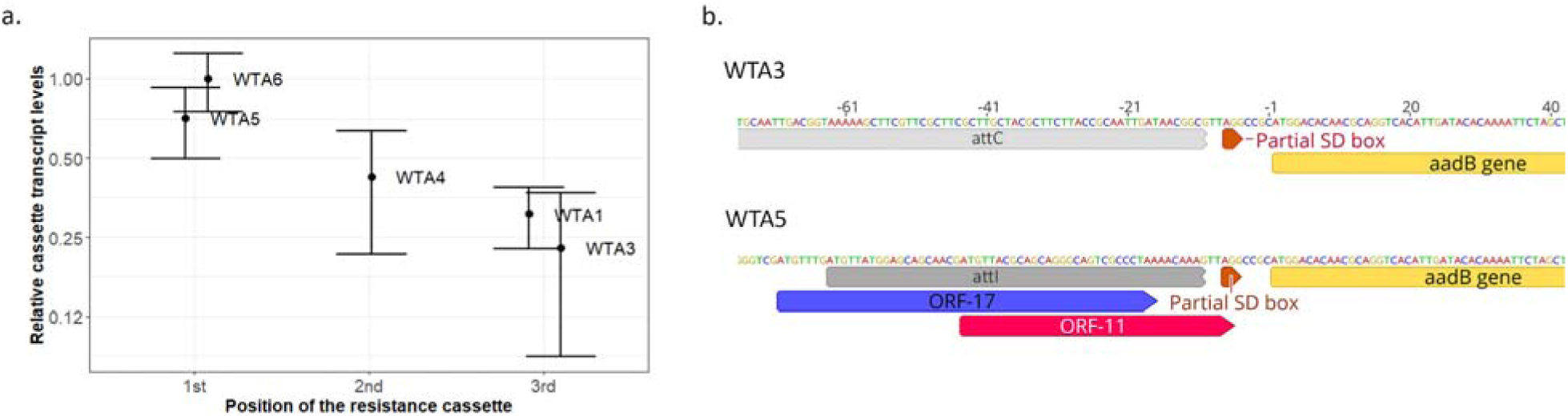
Transcriptional and translational origin of the aadB expression gradient. A. Transcript levels of the *aadB* cassette in the different arrays. Transcription levels are normalized relatively to the best transcribed array (WTA6). Error bars represent the standard error of three independent biological replicates. B. Representation of the genomic environment of the *aadB* cassette when *aadB* is in last (WTA3) or first position (WTA5) in our arrays. The two ORFs overlapping the *attI* which have been shown to improve cassette translation (Hanau-Berçot et al., 2002; Papagiannitsis et al., 2017) are represented in colour.

**Fig S2.**
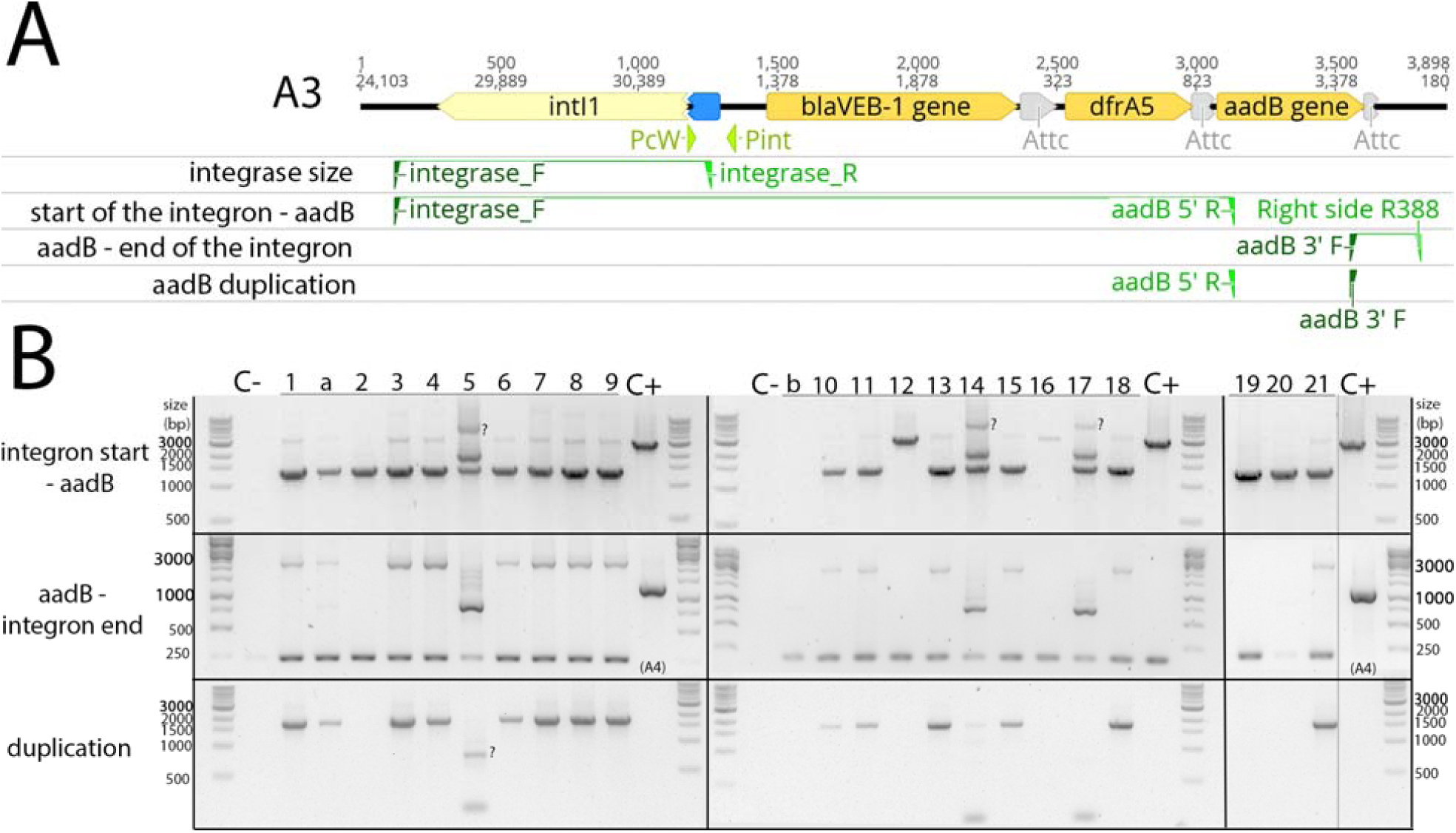
Rearrangements detection by PCR in x1024 MIC WTA3 populations: A. Primer binding sites and amplicons used in the screen for cassette rearrangements. B. Rearrangement PCR screen of the x1024 WTA3 populations. The population number identifier is indicated at the top. Samples with a letter were excluded from the analysis due to contamination. Bands which cannot be explained by the genomic data are indicated by a question mark. Non-relevant lanes were excluded from the gels on the right where indicated by the grey vertical lane and care was taken to keep the vertical alignment within the gels pictures during the figure composition.

**Figure S3:**
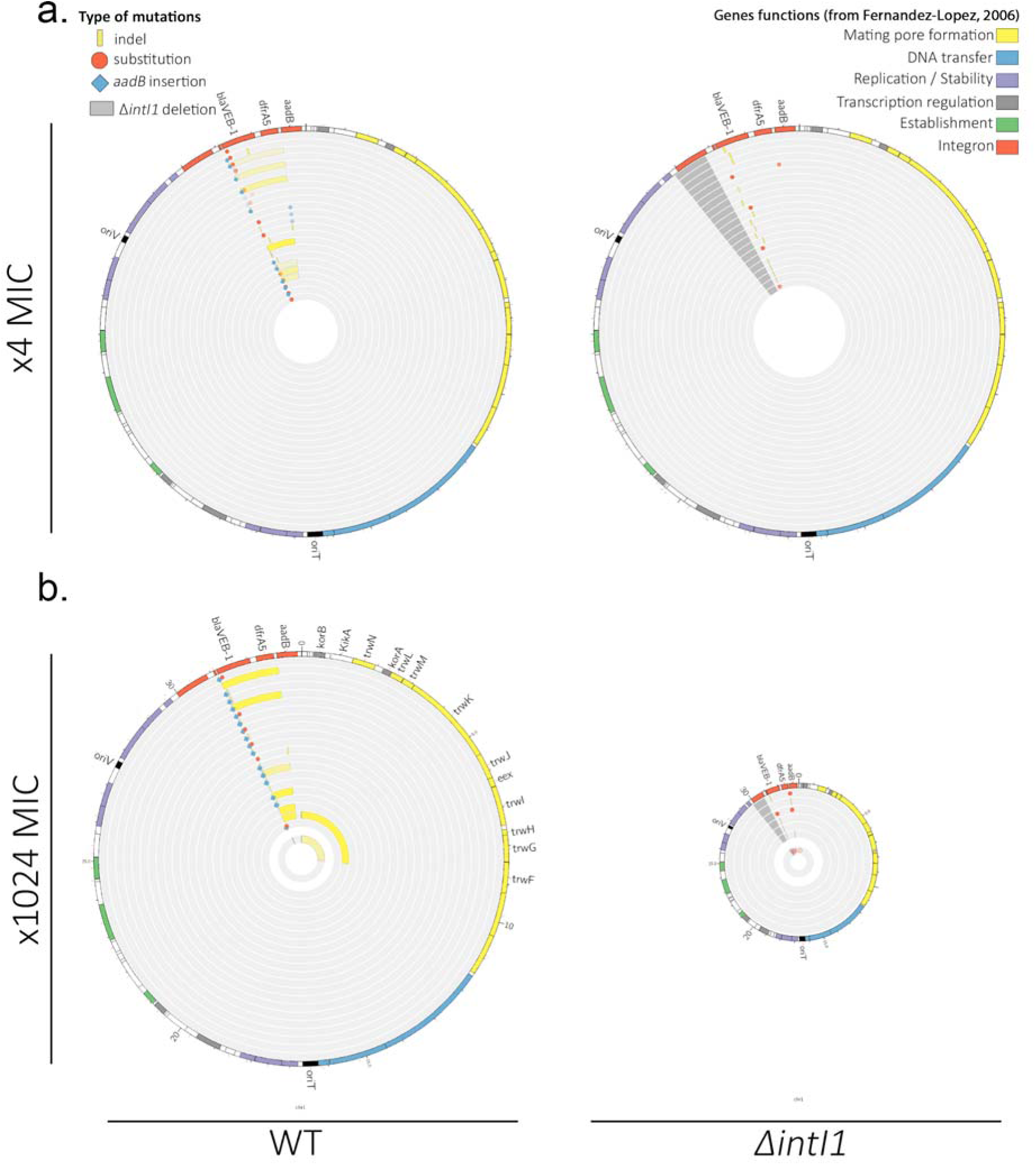
Plasmid mutations and rearrangements at (a) x4 MIC and (b) x1024 MIC: Representation of the plasmid mutations and rearrangements in PAO1:WTA3 and PAO1: Δ*intI1*A3 populations, mapped to the plasmid reference sequence. Each circle represents a separate population. Indels are represented in yellow and single nucleotide substitutions in red. aadB insertions are represented by blue lozenges. The color intensity represents the frequency of the corresponding mutation/recombination. The dark grey area in the PAO1:Δ*intI1* populations represents the location of the intI1 deletion. The function of each R388 gene as described in (Fernández-López et al., 2006) is indicated by a specific color in the outer circle. Apart from oriT and oriV, given as reference, only the name of genes where mutations are located is indicated. In (b) the inner circles correspond to the equimolar ⅛ MIC control populations.

**Fig S4.**
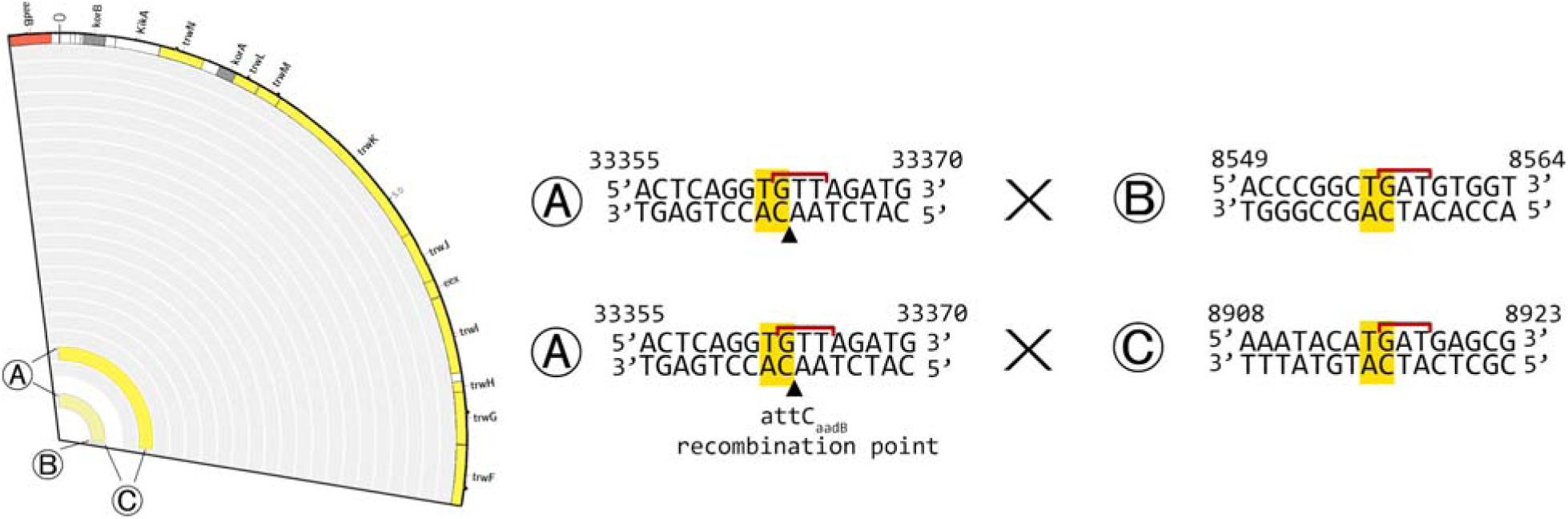
Left: Rearrangements in the WT populations at x1024 MIC. Each junction site is indicated by a letter. Right: junction sequences for each rearrangement. The junction site is indicated in yellow (as the crossover point is unclear due to sequence homology between each junction, the entire homology is highlighted). The GNT integron secondary motif is indicated by a red line. The recombination point of the *aadB* attC site is indicated by a black arrow.

**Fig S5.**
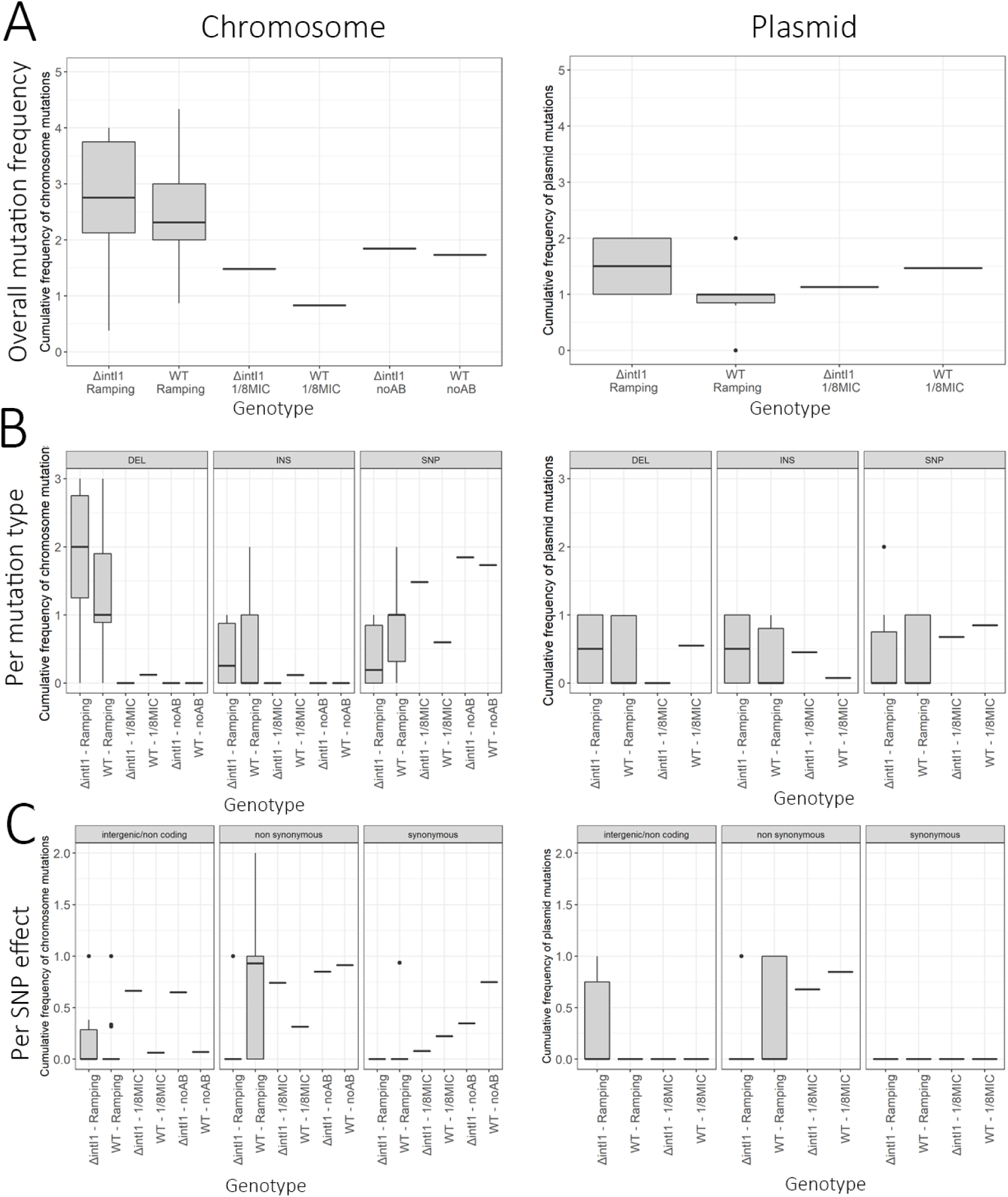
Summary statistics of mutations in the x1024 MIC populations. Box plots representing the average cumulative mutation frequencies for each population at the x1024 MIC time-point for A. all mutations B. per mutation type C. per mutation effect (SNP only). The lower and upper hinges correspond to the first and third quartiles while the middle line corresponds to the median. Cassette rearrangements (duplications and deletions) are not included

**Fig S6.**
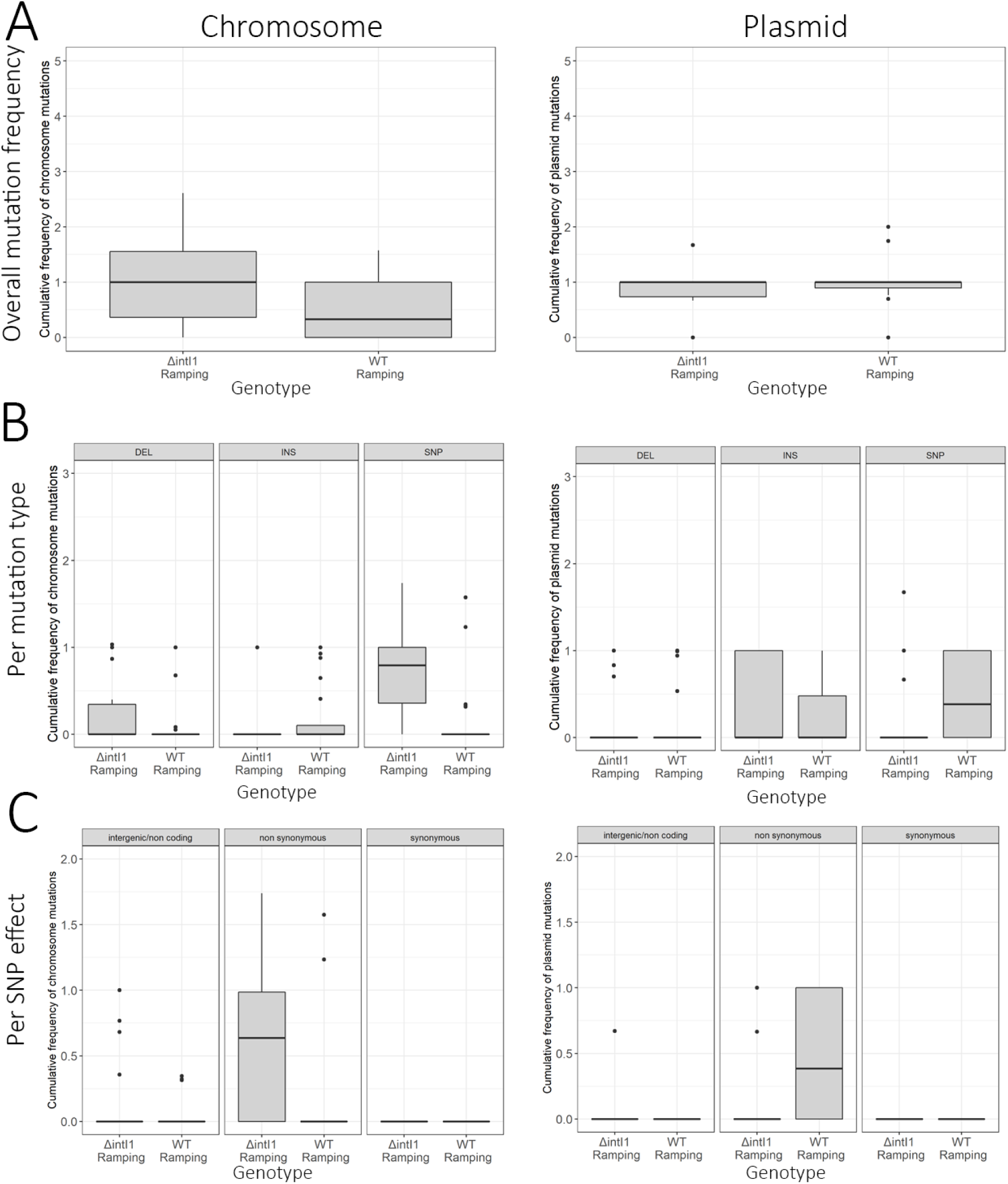
Summary statistics of mutations in the x4 MIC populations. Box plots representing the average cumulative mutation frequencies for each population at the x4 MIC time-point for A. all mutations B. per mutation type C. per mutation effect (SNP only). The lower and upper hinges correspond to the first and third quartiles while the middle line corresponds to the median. Cassette rearrangements (duplications and deletions) are not included.

**Fig S7.**
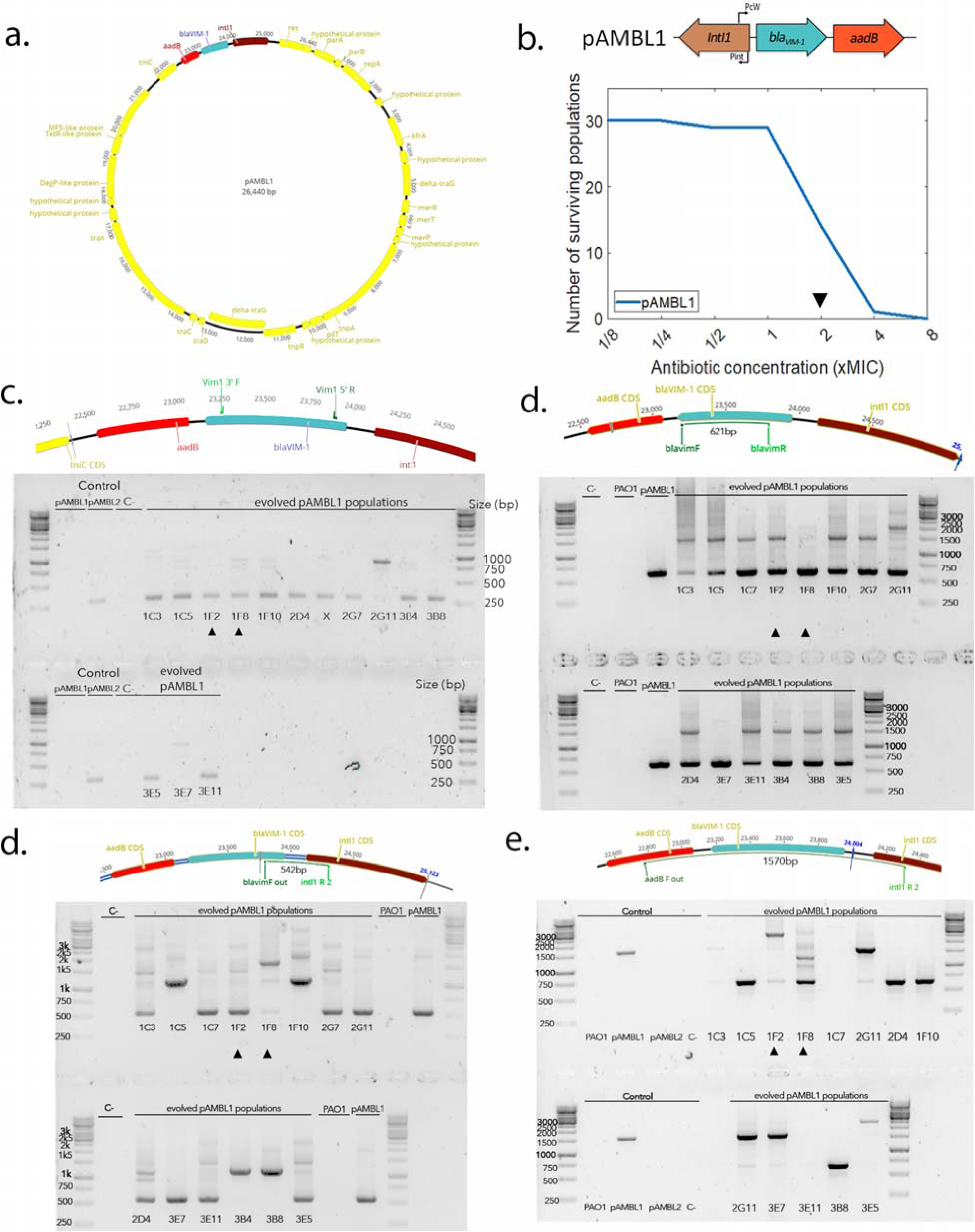
pAMBL1 rearrangements. a. Representation of the pAMBL1 plasmid. The integron is highlighted in color. b. Survival curve of the 30 PA01:pAMBL1 populations under ramping treatment. The black arrow indicates the time-point at which populations were plated and duplications amplified by PCR/NGS. c.to e. Detection of cassettes rearrangements by PCR. The expected positions of the primers on the ancestral pAMBL1 and the size of the corresponding amplicon is indicated on top.

**Figure. S8:**
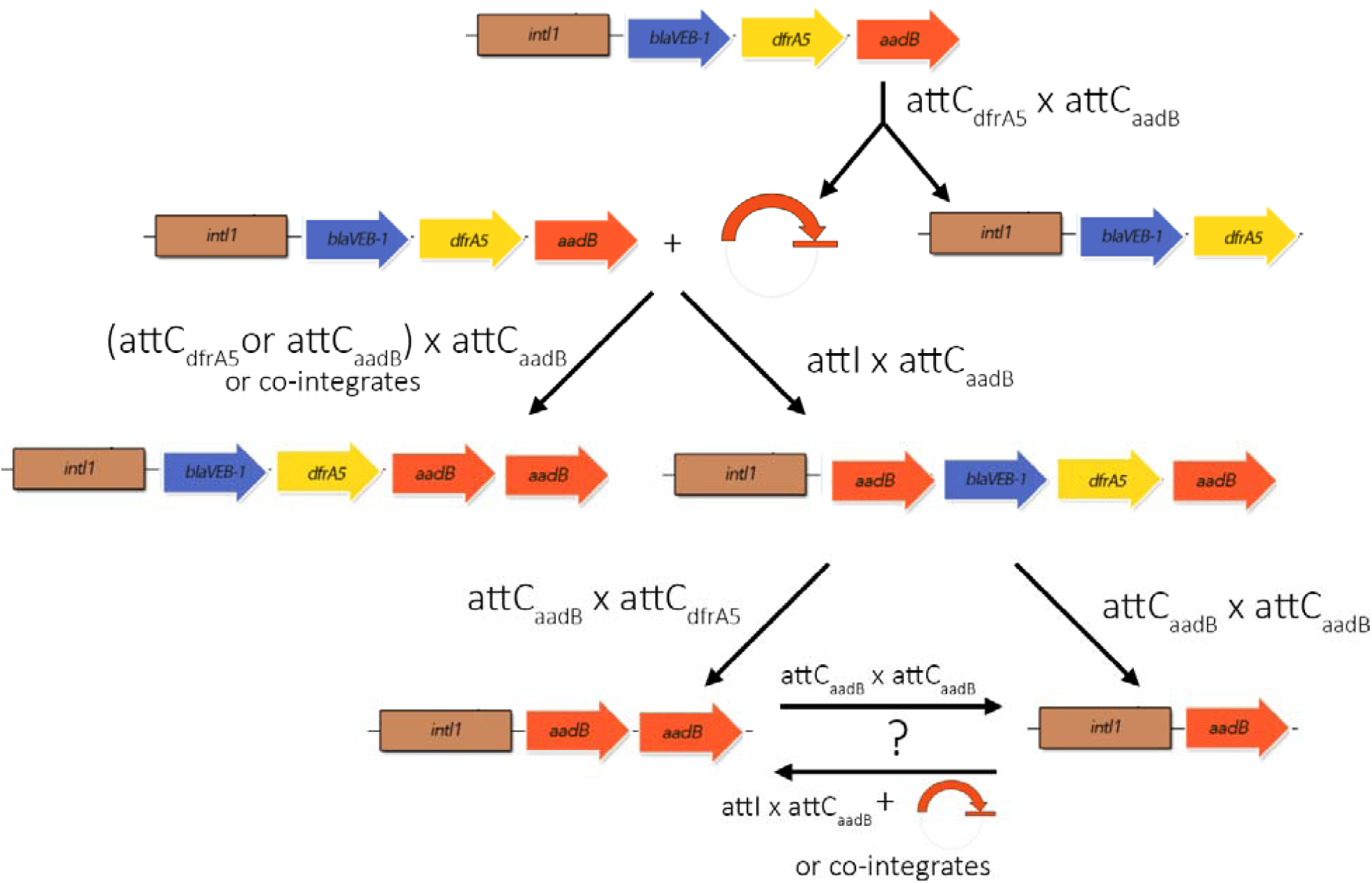
Representation of the proposed reactions leading to the various arrays observed in this study.

## Supplementary Tables

**Table S1 – Strains used in this study**

**Table S2 – Primers used in this study**

**Table S3 – List of mutations / recombinations at x1024 MIC**

**Table S4 – List of mutations / recombinations at x4 MIC**

**Table S5 – List of duplicated cassettes from the INTEGRALL database**

## Notes

### Competing Interest Statement

The authors have declared no competing interest.

### Summary of Updates

Slight update of the abstract and addition of accession number for genomic data and Dryad data repository doi

## Bibliography

Alkan, C., Sajjadian, S., & Eichler, E. E. (2011). Limitations of next-generation genome sequence assembly. Nature Methods, 8(1), 61–65. https://doi.org/10.1038/nmeth.1527

Andersson, D. I., & Hughes, D. (2009). Gene Amplification and Adaptive Evolution in Bacteria. Annual Review of Genetics, 43(1), 167–195. https://doi.org/10.1146/annurev-genet-102108-134805

Antipov, D., Hartwick, N., Shen, M., Raiko, M., Lapidus, A., & Pevzner, P. A. (2016). plasmidSPAdes: Assembling plasmids from whole genome sequencing data. Bioinformatics, 32(22), btw493–btw493. https://doi.org/10.1093/bioinformatics/btw493

Aubert, D., Naas, T., & Nordmann, P. (2012). Integrase-mediated recombination of the veb1 gene cassette encoding an extended-spectrum β-lactamase. PloS One, 7(12), e51602–e51602. https://doi.org/10.1371/journal.pone.0051602

Avila, P., & de la Cruz, F. (1988). Physical and genetic map of the IncW plasmid R388. Plasmid, 20(2), 155–157. https://doi.org/10.1016/0147-619X(88)90019-4

Bankevich, A., Nurk, S., Antipov, D., Gurevich, A. A., Dvorkin, M., Kulikov, A. S., Lesin, V. M., Nikolenko, S. I., Pham, S., Prjibelski, A. D., Pyshkin, A. V., Sirotkin, A. V., Vyahhi, N., Tesler, G., Alekseyev, M. A., & Pevzner, P. A. (2012). SPAdes: A New Genome Assembly Algorithm and Its Applications to Single-Cell Sequencing. Journal of Computational Biology, 19(5), 455–477. https://doi.org/10.1089/cmb.2012.0021

Barraud, O., & Ploy, M.-C. (2015). Diversity of Class 1 Integron Gene Cassette Rearrangements Selected under Antibiotic Pressure. Journal of Bacteriology, 197(13), 2171–2178. https://doi.org/10.1128/JB.02455-14

Barrick, J. E., Colburn, G., Deatherage, D. E., Traverse, C. C., Strand, M. D., Borges, J. J., Knoester, D. B., Reba, A., & Meyer, A. G. (2014). Identifying structural variation in haploid microbial genomes from short-read resequencing data using breseq. BMC Genomics, 15(1), 1039–1039. https://doi.org/10.1186/1471-2164-15-1039

Bolger, A. M., Lohse, M., & Usadel, B. (2014). Trimmomatic: A flexible trimmer for Illumina sequence data. Bioinformatics (Oxford, England), 30(15), 2114–2120. https://doi.org/10.1093/bioinformatics/btu170

Brook, I. (1989). Inoculum Effect. Reviews of Infectious Diseases, 11(3), 361–368.

Brynildsrud, O. (2018). Read Depth Analysis to Identify CNV in Bacteria Using CNOGpro (pp. 73–81). Humana Press, New York, NY. https://doi.org/10.1007/978-1-4939-8666-8_5

Cambray, G., Sanchez-Alberola, N., Campoy, S., Guerin, É., Da Re, S., González-Zorn, B., Ploy, M.-C., Barbé, J., Mazel, D., Erill, I., Stokes, H., Hall, R., Collis, C., Kim, M., Stokes, H., Hall, R., Levesque, C., Brassard, S., Lapointe, J., … Brenner, S. (2011). Prevalence of SOS-mediated control of integron integrase expression as an adaptive trait of chromosomal and mobile integrons. Mobile DNA, 2(1), 6–6. https://doi.org/10.1186/1759-8753-2-6

Cameron, F. H., Groot Obbink, D. J., Ackerman, V. P., & Hall, R. M. (1986). Nucleotide sequence of the AAD(2′) aminoglycoside adenylyltransferase determinant aadB. Evolutionary relationship of this region with those surrounding aadA in R538-1 and dhfrll in R388. Nucleic Acids Research, 14(21), 8625–8635. https://doi.org/10.1093/nar/14.21.8625

Chen, J., Li, H., Yang, J., Zhan, R., Chen, A., & Yan, Y. (2015). Prevalence and Characterization of Integrons in Multidrug Resistant Acinetobacter baumannii in Eastern China: A Multiple-Hospital Study. International Journal of Environmental Research and Public Health, 12(8), 10093–10105. https://doi.org/10.3390/ijerph120810093

Collis, C. M., & Hall, R. M. (1992). Site-specific deletion and rearrangement of integron insert genes catalyzed by the integron DNA integrase. Journal of Bacteriology, 174(5), 1574–1585.

Collis, C. M., & Hall, R. M. (1995). Expression of antibiotic resistance genes in the integrated cassettes of integrons. Antimicrobial Agents and Chemotherapy, 39(1), 155–162.

Dettman, J. R., Rodrigue, N., & Kassen, R. (2015). Genome-Wide Patterns of Recombination in the Opportunistic Human Pathogen Pseudomonas aeruginosa. Genome Biology and Evolution, 7(1), 18–34. https://doi.org/10.1093/gbe/evu260

Engelstädter, J., Harms, K., & Johnsen, P. J. (2016). The evolutionary dynamics of integrons in changing environments. The ISME Journal. https://doi.org/10.1038/ismej.2015.222

Escudero, J. A., Loot, C., Nivina, A., & Mazel, D. (2015). The Integron: Adaptation On Demand. Microbiology Spectrum, 3(2), MDNA3–0019–2014. https://doi.org/10.1128/microbiolspec.MDNA3-0019-2014

Ewels, P., Magnusson, M., Lundin, S., & Käller, M. (2016). MultiQC: summarize analysis results for multiple tools and samples in a single report. Bioinformatics, 32(19), 3047–3048. https://doi.org/10.1093/bioinformatics/btw354

Fernández-López, R., Garcillán-Barcia, M. P., Revilla, C., Lázaro, M., Vielva, L., & de la Cruz, F. (2006). Dynamics of the IncW genetic backbone imply general trends in conjugative plasmid evolution. FEMS Microbiology Reviews, 30(6), 942–966. https://doi.org/10.1111/j.1574-6976.2006.00042.x

Firoozeh, F., Mahluji, Z., Khorshidi, A., & Zibaei, M. (2019). Molecular characterization of class 1, 2 and 3 integrons in clinical multi-drug resistant Klebsiella pneumoniae isolates. Antimicrobial Resistance & Infection Control, 8(1), 59. https://doi.org/10.1186/s13756-019-0509-3

Foster, P. L. (2011). Stress-induced Mutagenesis in Bacteria. In ELS (pp. 89–104). John Wiley & Sons, Ltd. https://doi.org/10.1002/9780470015902.a0023608

Gassama, A., Aídara-Kane, A., Chainier, D., Denis, F., & Ploy, M. C. (2004). Integron-Associated Antibiotic Resistance in Enteroaggregative and Enteroinvasive Escherichia coli. Microbial Drug Resistance, 10(1), 27–30. https://doi.org/10.1089/107662904323047763

Gifford, D., Furió, V., Papkou, A., Vogwill, T., Oliver, A., & MacLean, R. C. (2018). Identifying and exploiting genes that potentiate the evolution of antibiotic resistance. Nature Ecology & Evolution, 2(6), 1033–1039. https://doi.org/10.1038/s41559-018-0547-x

Guerin, E., Cambray, G., Sanchez-Alberola, N., Campoy, S., Erill, I., Da Re, S., Gonzalez-Zorn, B., Barbé, J., Ploy, M.-C., & Mazel, D. (2009). The SOS response controls integron recombination. Science (New York, N.Y.), 324 (5930), 1034–1034. https://doi.org/10.1126/science.1172914

Halaji, M., Feizi, A., Mirzaei, A., Sedigh Ebrahim-Saraie, H., Fayyazi, A., Ashraf, A., & Havaei, S. A. (2020). The Global Prevalence of Class 1 Integron and Associated Antibiotic Resistance in Escherichia coli from Patients with Urinary Tract Infections, a Systematic Review and Meta-Analysis. Microbial Drug Resistance. https://doi.org/10.1089/mdr.2019.0467

Hall, R M, Brookes, D. E., & Stokes, H. W. (1991). Site-specific insertion of genes into integrons: Role of the 59-base element and determination of the recombination cross-over point. Molecular Microbiology, 5(8), 1941–1959.

Hall, Ruth M., Recchia, G. D., Scaramuzzi, C., Stokes, H. W., Partridge, S. R., & Collis, C. M. (2000). Definition of the attI1 site of class 1 integrons. Microbiology, 146(11), 2855–2864. https://doi.org/10.1099/00221287-146-11-2855

Hanau-Berçot, B., Podglajen, I., Casin, I., & Collatz, E. (2002). An intrinsic control element for translational initiation in class 1 integrons. Molecular Microbiology, 44(1), 119–130. https://doi.org/10.1046/j.1365-2958.2002.02843.x

Harms, K., Starikova, I., & Johnsen, P. J. (2013). Costly Class-1 integrons and the domestication of the the functional integrase. Mobile Genetic Elements, 3(2), e24774–e24774. https://doi.org/10.4161/mge.24774

Higgins, S., Heeb, S., Rampioni, G., Fletcher, M. P., Williams, P., & Cámara, M. (2018). Differential Regulation of the Phenazine Biosynthetic Operons by Quorum Sensing in Pseudomonas aeruginosa PAO1-N. Frontiers in Cellular and Infection Microbiology, 8, 252–252. https://doi.org/10.3389/fcimb.2018.00252

Hocquet, D., Llanes, C., Thouverez, M., & al., et. (2012). Evidence for Induction of Integron-Based Antibiotic Resistance by the SOS Response in a Clinical Setting. PLoS Pathogens, 8(6), e1002778–e1002778. https://doi.org/10.1371/journal.ppat.1002778

Jacquier, H., Zaoui, C., Sanson-le Pors, M.-J., Mazel, D., & Berçot, B. (2009). Translation regulation of integrons gene cassette expression by the attC sites. Molecular Microbiology, 72(6), 1475–1486. https://doi.org/10.1111/j.1365-2958.2009.06736.x

Jové, T., Da Re, S., Denis, F., Mazel, D., & Ploy, M.-C. (2010). Inverse correlation between promoter strength and excision activity in class 1 integrons. PLoS Genetics, 6(1), e1000793–e1000793. https://doi.org/10.1371/journal.pgen.1000793

Kishony, R., & Leibler, S. (2003). Environmental stresses can alleviate the average deleterious effect of mutations. Journal of Biology, 2(2). https://doi.org/10.1186/1475-4924-2-14

Kohanski, M. A., Dwyer, D. J., & Collins, J. J. (2010). How antibiotics kill bacteria: From targets to networks. Nature Reviews Microbiology, 8(6), 423–435. https://doi.org/10.1038/nrmicro2333

Krzywinski, M., Schein, J., Birol, I., Connors, J., Gascoyne, R., Horsman, D., Jones, S. J., & Marra, M. A. (2009). Circos: An information aesthetic for comparative genomics. Genome Research, 19(9), 1639–1645. https://doi.org/10.1101/gr.092759.109

Lacotte, Y., Ploy, M.-C., & Raherison, S. (2017). Class 1 integrons are low-cost structures in Escherichia coli. The ISME Journal, 11(7), 1535–1544. https://doi.org/10.1038/ismej.2017.38

Lau, C. H.-F., Fraud, S., Jones, M., Peterson, S. N., & Poole, K. (2013). Mutational activation of the AmgRS two-component system in aminoglycoside-resistant Pseudomonas aeruginosa. Antimicrobial Agents and Chemotherapy, 57(5), 2243–2251. https://doi.org/10.1128/AAC.00170-13

Lau, C. H.-F., Krahn, T., Gilmour, C., Mullen, E., & Poole, K. (2015). AmgRS-mediated envelope stress-inducible expression of the mexXY multidrug efflux operon of Pseudomonas aeruginosa. MicrobiologyOpen, 4(1), 121–135. https://doi.org/10.1002/mbo3.226

Levesque, R. C. (2006). In Vivo Functional Genomics of Pseudomonas: PCR-Based Signature-Tagged Mutagenesis. In Pseudomonas (pp. 99–120). Springer US. https://doi.org/10.1007/0-387-28881-3_4

Li, B., Hu, Y., Wang, Q., Yi, Y., Woo, P. C. Y., Jing, H., Zhu, B., & Liu, C. H. (2013). Structural Diversity of Class 1 Integrons and Their Associated Gene Cassettes in Klebsiella pneumoniae Isolates from a Hospital in China. PLOS ONE, 8(9), e75805. https://doi.org/10.1371/journal.pone.0075805

Liebert, C. A., Hall, R. M., & Summers, A. O. (1999). Transposon Tn21, flagship of the floating genome. Microbiology and Molecular Biology Reviews?: MMBR, 63(3), 507–522.

Loot, C., Ducos-Galand, M., Escudero, J. A., Bouvier, M., & Mazel, D. (2012). Replicative resolution of integron cassette insertion. Nucleic Acids Research, 40(17), 8361–8370. https://doi.org/10.1093/nar/gks620

López-Causapé, C., Cabot, G., del Barrio-Tofiño, E., & Oliver, A. (2018). The Versatile Mutational Resistome of Pseudomonas aeruginosa. Frontiers in Microbiology, 9. https://doi.org/10.3389/fmicb.2018.00685

MacLean, R. C., Torres-Barceló, C., & Moxon, R. (2013). Evaluating evolutionary models of stress-induced mutagenesis in bacteria. Nature Reviews Genetics, 14(3), 221–227. https://doi.org/10.1038/nrg3415

Mitsuhashi, S., Harada, K., Hashimoto, H., & Egawa, R. (1961). On the drug-resistance of enteric bacteria. 4. Drug-resistance of Shigella prevalent in Japan. The Japanese Journal of Experimental Medicine, 31, 47–52.

Moura, A., Soares, M., Pereira, C., Leitño, N., Henriques, I., & Correia, A. (2009). INTEGRALL: A database and search engine for integrons, integrases and gene cassettes. Bioinformatics, 25(8), 1096–1098. https://doi.org/10.1093/bioinformatics/btp105

Okii, M., Iyobe, S., & Mitsuhashi, S. (1983). Mapping of the gene specifying aminoglycoside 3’-phosphotransferase II on the Pseudomonas aeruginosa chromosome. Journal of Bacteriology, 155(2), 643–649.

Oliver, A., Mulet, X., López-Causapé, C., & Juan, C. (2015). The increasing threat of Pseudomonas aeruginosa high-risk clones. Drug Resistance Updates, 21–22, 41–59. https://doi.org/10.1016/j.drup.2015.08.002

Papagiannitsis, C. C., Tzouvelekis, L. S., Tzelepi, E., & Miriagou, V. (2017). AttI1-Located small open reading frames ORF-17 and ORF-11 in a class 1 integron affect expression of a gene cassette possessing a canonical shine-dalgarno sequence. Antimicrobial Agents and Chemotherapy, 61(3). https://doi.org/10.1128/AAC.02070-16

Partridge, S. R., Kwong, S. M., Firth, N., & Jensen, S. O. (2018). Mobile Genetic Elements Associated with Antimicrobial Resistance. Clinical Microbiology Reviews, 31(4). https://doi.org/10.1128/CMR.00088-17

Picard toolkit. (n.d.). Broad Institute. http://broadinstitute.github.io/picard/

Poirel, L., Naas, T., Guibert, M., Chaibi, E. B., Labia, R., & Nordmann, P. (1999). Molecular and biochemical characterization of VEB-1, a novel class A extended-spectrum beta-lactamase encoded by an Escherichia coli integron gene. Antimicrobial Agents and Chemotherapy, 43(3), 573–581.

Providenti, M. A., O’Brien, J. M., Ewing, R. J., Paterson, E. S., & Smith, M. L. (2006). The copy-number of plasmids and other genetic elements can be determined by SYBR-Green-based quantitative real-time PCR. Journal of Microbiological Methods, 65(3), 476–487. https://doi.org/10.1016/J.MIMET.2005.09.007

Rao, A. N., Barlow, M., Clark, L. A., Boring, J. R., Tenover, F. C., & McGowan, J. E. (2006). Class 1 integrons in resistant Escherichia coli and Klebsiella spp., US hospitals. Emerging Infectious Diseases, 12(6), 1011–1014. https://doi.org/10.3201/eid1206.051596

Recchia, G. D., & Hall, R. M. (1995). Gene cassettes: A new class of mobile element. Microbiology, 141(12), 3015–3027. https://doi.org/10.1099/13500872-141-12-3015

Recchia, G. D., Stokes, H. W., & Hall, R. M. (1994). Characterisation of specific and secondary recombination sites recognised by the integron DNA integrase. Nucleic Acids Research, 22(11), 2071–2078. https://doi.org/10.1093/nar/22.11.2071

Rownd, R., Nakaya, R., & Nakamura, A. (1966). Molecular nature of the drug-resistance factors of the enterobacteriaceae. Journal of Molecular Biology, 17(2), 376–393. https://doi.org/10.1016/S0022-2836(66)80149-3

Ruiz-Martínez, L., López-Jiménez, L., Fusté, E., Vinuesa, T., Martínez, J. P., & Viñas, M. (2011). Class 1 integrons in environmental and clinical isolates of Pseudomonas aeruginosa. International Journal of Antimicrobial Agents, 38(5), 398–402. https://doi.org/10.1016/j.ijantimicag.2011.06.016

San Millan, A., Escudero, J. A., Gifford, D. R., Mazel, D., & MacLean, R. C. (2016). Multicopy plasmids potentiate the evolution of antibiotic resistance in bacteria. Nature Ecology & Evolution, 1(1), 1–8. https://doi.org/10.1038/s41559-016-0010

San Millan, A., Santos-Lopez, A., Ortega-Huedo, R., Bernabe-Balas, C., Kennedy, S. P., & Gonzalez-Zorn, B. (2015). Small-plasmid-mediated antibiotic resistance is enhanced by increases in plasmid copy number and bacterial fitness. Antimicrobial Agents and Chemotherapy, 59(6), 3335–3341. https://doi.org/10.1128/AAC.00235-15

San Millan, A., Toll-Riera, M., Escudero, J. A., Cantón, R., Coque, T. M., & Craig MacLean, R. (2015). Sequencing of plasmids pAMBL1 and pAMBL2 from Pseudomonas aeruginosa reveals a blaVIM-1 amplification causing high-level carbapenem resistance. Journal of Antimicrobial Chemotherapy, 70(11), 3000–3003. https://doi.org/10.1093/jac/dkv222

Schurek, K. N., Marr, A. K., Taylor, P. K., Wiegand, I., Semenec, L., Khaira, B. K., & Hancock, R. E. W. (2008). Novel genetic determinants of low-level aminoglycoside resistance in Pseudomonas aeruginosa. Antimicrobial Agents and Chemotherapy, 52(12), 4213–4219. https://doi.org/10.1128/AAC.00507-08

Seemann, T. (2014). Prokka: Rapid prokaryotic genome annotation. Bioinformatics, 30(14), 2068–2069. https://doi.org/10.1093/bioinformatics/btu153

Simon Andrews. (n.d.). FastQC. FastQC: A Quality Control Tool for High Throughput Sequence Data [Online]. http://www.bioinformatics.babraham.ac.uk/projects/fastqc/

Starikova, I., Harms, K., Haugen, P., Lunde, T. T. M., Primicerio, R., Samuelsen, Ø., Nielsen, K. M., & Johnsen, P. J. (2012). A trade-off between the fitness cost of functional integrases and long-term stability of integrons. PLoS Pathogens, 8(11), e1003043–e1003043. https://doi.org/10.1371/journal.ppat.1003043

Stokes, H. W., & Hall, R. M. (1989). A novel family of potentially mobile DNA elements encoding site-specific gene-integration functions: Integrons. Molecular Microbiology, 3(12), 1669–1683.

Stokes, H. W., & Hall, R. M. (1992). The integron In1 in plasmid R46 includes two copies of the oxa2 gene cassette. Plasmid, 28(3), 225–234. https://doi.org/10.1016/0147-619X(92)90054-E

Sundström, L., Rådström, P., Swedberg, G., & Sköld, O. (1988). Site-specific recombination promotes linkage between trimethoprim- and sulfonamide resistance genes. Sequence characterization of dhfrV and sulI and a recombination active locus of Tn 21. MGG Molecular & General Genetics, 213 (2–3), 191–201. https://doi.org/10.1007/BF00339581

Therneau, T. M. (2020). A Package for Survival Analysis in R. https://CRAN.R-project.org/package=survival

Torres-Barceló, C., Kojadinovic, M., Moxon, R., MacLean, R. C., Goh, EB., Yim, G., Tsui, W., McClure, J., Surette, MG., Davies, J., Khil, PP., Camerini-Otero, RD., Baharoglu, Z., Mazel, D., MacLean, RC., Torres-Barceló, C., Moxon, R., Shee, C., Gibson, JL., … Collins, JJ. (2015). The SOS response increases bacterial fitness, but not evolvability, under a sublethal dose of antibiotic. Proceedings. Biological Sciences / The Royal Society, 282 (1816), 20150885–20150885. https://doi.org/10.1098/rspb.2015.0885

Turton, J. F., Kaufmann, M. E., Glover, J., Coelho, J. M., Warner, M., Pike, R., & Pitt, T. L. (2005). Detection and Typing of Integrons in Epidemic Strains of Acinetobacter baumannii Found in the United Kingdom. Journal of Clinical Microbiology, 43(7), 3074–3082. https://doi.org/10.1128/JCM.43.7.3074-3082.2005

Vogne, C., Aires, J. R., Bailly, C., Hocquet, D., & Plésiat, P. (2004). Role of the multidrug efflux system MexXY in the emergence of moderate resistance to aminoglycosides among Pseudomonas aeruginosa isolates from patients with cystic fibrosis. Antimicrobial Agents and Chemotherapy, 48(5), 1676–1680. https://doi.org/10.1128/aac.48.5.1676-1680.2004

Wick, R. R., Schultz, M. B., Zobel, J., & Holt, K. E. (2015). Bandage: Interactive visualization of de novo genome assemblies. Bioinformatics, 31(20), 3350–3352. https://doi.org/10.1093/bioinformatics/btv383

Yu, H. S., Lee, J. C., Kang, H. Y., Ro, D. W., Chung, J. Y., Jeong, Y. S., Tae, S. H., Choi, C. H., Lee, E. Y., Seol, S. Y., Lee, Y. C., & Cho, D. T. (2003). Changes in gene cassettes of class 1 integrons among Escherichia coli isolates from urine specimens collected in Korea during the last two decades. Journal of Clinical Microbiology, 41(12), 5429–5433. https://doi.org/10.1128/jcm.41.12.5429-5433.2003

